# Structure and conformational dynamics of the *Pseudomonas* CbrA transceptor

**DOI:** 10.64898/2026.03.10.710862

**Authors:** Melanie A. Orlando, Tejas Shah, Matthew W. Faber, Samik Bose, Benjamin J. Orlando

**Affiliations:** Department of Biochemistry and Molecular Biology, Michigan State University, East Lansing, MI 48824; Department of Computational Mathematics, Science and Engineering, Michigan State University, East Lansing, Michigan 48824, United States

**Keywords:** CbrA, transceptor, SLC transporter, molecular dynamics

## Abstract

The CbrA protein is a central regulator of carbon metabolism, biofilm formation, and virulence in *Pseudomonas* species, but the molecular mechanisms by which CbrA links nutrient sensing to downstream signaling has remained unclear. CbrA is a rare “transceptor” that combines membrane transporter and histidine kinase domains into a single functional polypeptide. The structural basis for histidine recognition and membrane transport, as well as signaling through intracellular histidine kinase domains has remained elusive. Here we determined a cryo-EM structure of CbrA which provides key molecular details of the SLC5-STAC domains in this unusual system. Unexpectedly, the small peptide CbrX encoded upstream of CbrA formed a stable complex with the SLC5 transporter domain. The structure reveals how histidine binds within the transporter, and molecular dynamics simulations provide insight into proton gradient driven conformational changes that enable histidine transport. These findings define the molecular architecture of key CbrA functional domains, and pave a path toward developing a comprehensive understanding of coupling between membrane transport and downstream signaling pathways that guide essential physiological traits in *Pseudomonas*.

**Significance Statement:** CbrA is a key regulator of carbon–nitrogen metabolism in *Pseudomonas* and is essential for successful host infection. The molecular basis for CbrA’s dual role in membrane transport and downstream signaling has remained elusive. Here we determined a cryo-EM structure that defines the organization of the CbrA SLC5 and STAC domains, and reveals that the small peptide CbrX encoded upstream of CbrA forms a stable complex with the SLC5 transporter region. A structure with histidine trapped in the binding cavity, together with molecular dynamics simulations, identifies protonation dependent transitions that guide the transport cycle. This work establishes a mechanistic foundation for understanding how CbrA and related transceptors integrate substrate sensing and transport with regulatory control of signaling.

## Introduction

*Pseudomonadaceae* are metabolically versatile bacteria that inhabit diverse environmental niches, with *Pseudomonas aeruginosa* in particular acting as an opportunistic pathogen that can cause serious life-threatening infections(1, 2). The ecological fitness and clinical significance of *P. aeruginosa* is underpinned by the organism’s ability to rapidly adapt to varying nutrient conditions(3), form biofilms(4), and increasingly develop resistance to multiple antimicrobial agents(1, 2). A central mediator in the adaptability of *Pseudomonaceae* is the CbrA/B regulatory network(5). The CbrA/B system integrates environmental cues to coordinate carbon and nitrogen metabolism(6–8), biofilm formation(9), and other cellular processes critical for survival and virulence(10, 11). CbrA functions as a global regulatory hub by coordinating the detection of environmental cues like nutrient availability with phosphorylation and activation of the σ^54^-dependent transcription factor CbrB (**Figure 1A**). Activated CbrB has been shown to directly control the expression of at least 61 different genes(12), including those involved in amino acid biosynthesis and metabolism such as the histidine utilization (*hut*) operon (**Figure 1A**)(7, 8, 13, 14). In addition, activated CbrB has been shown to regulate production of the iron scavengers pyoverdine and pyochelin(11), as well as the small RNAs *crcZ* and *crcY* which sequester the Crc-Hfq complex to modulate carbon catabolite repression (**Figure 1A**)(15–18). Disruption of CbrA markedly alters *Pseudomonas* growth and biofilm formation under certain conditions(9), and significantly attenuates virulence and microbial burden in murine infection models(10). Given its central role in *Pseudomonas* physiology and pathogenicity, the CbrA/B network represents a compelling target for future therapeutic development.

**Figure 1.**
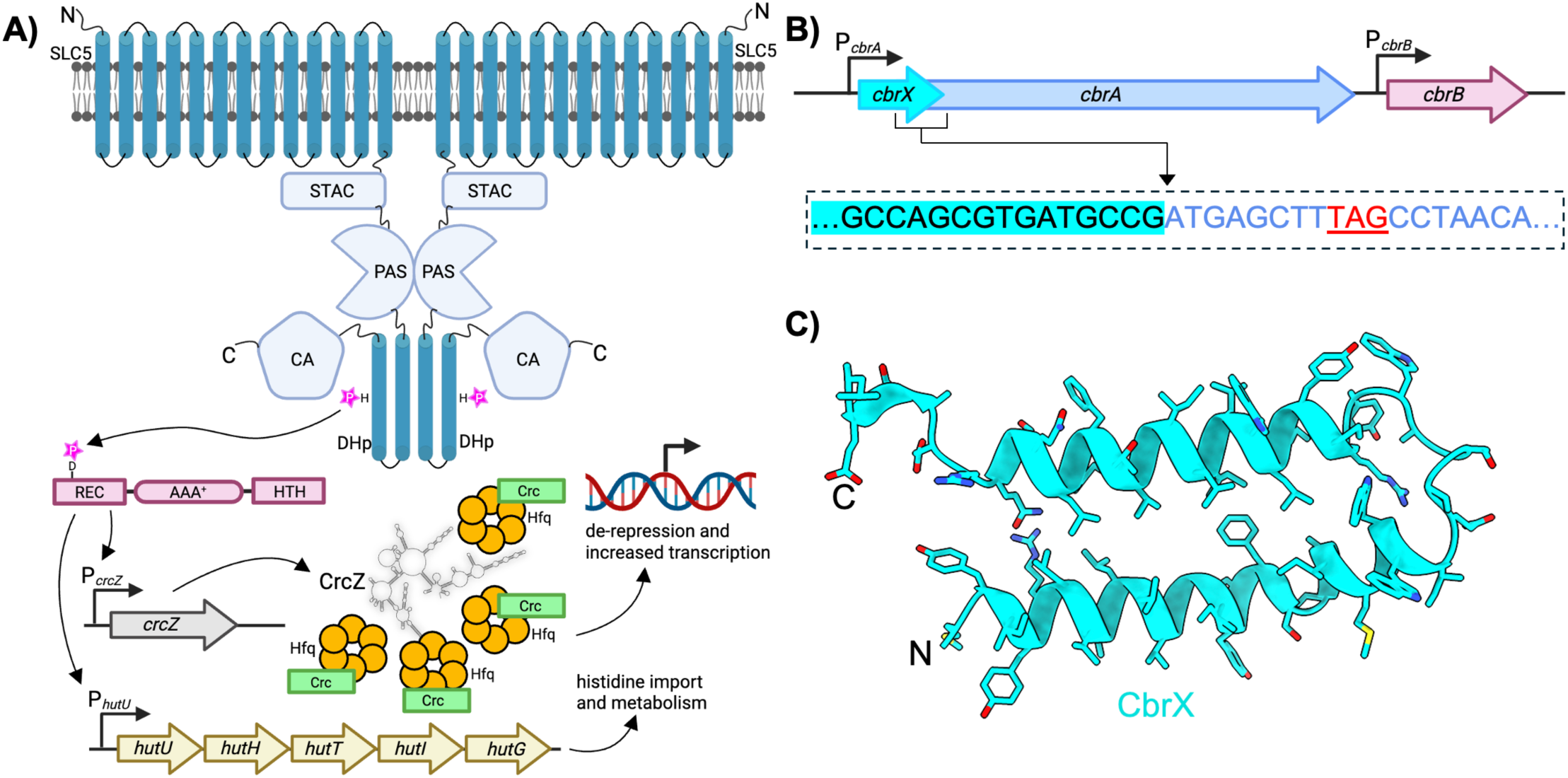
Topology and genomic organization of CbrXA. **A)** Schematic showing the overall domain organization of CbrA (blue) and CbrB (red). CbrA is composed of an N-terminal SLC5 transporter domain, followed by cytosolic STAC, PAS, DHp, and catalytic (CA) domains. The schematic shows CbrA as a homodimer, based on classical models of histidine kinase structure/function. CbrB (red) is composed of receiver (REC), AAA^+^ ATPase, and helix-turn-helix (HTH) subdomains. Phosphoryl group transfer between a conserved histidine in the DHp domain of CbrA and a conserved aspartate in the REC domain of CbrB serves to activate CbrB as a σ-54 transcription factor. **B)** Genomic organization of *cbrXA – cbrB* in *P. putida* KT2440. *CbrX* and *cbrA* are translationally coupled behind the *cbrA* promoter. The response regulator *cbrB* is under control of an independent promoter. The inset in the dashed outline shows genomic overlap of *cbrX* (cyan highlight, black letters) and *cbrA* (blue letters). The stop codon for *cbrX* is shown in red underlined letters. **C)** AlphaFold predicted secondary structure of cbrX, showing two alpha helices separated by a flexible linker.

CbrA is a rare example of a “transceptor” protein that combines membrane transporter and histidine kinase domains into a single polypeptide (**Figure 1A**). The N-terminus of CbrA consists of a solute carrier family 5 (SLC5) domain with 13 transmembrane (TM) helices embedded within the inner membrane. Previous studies have demonstrated that the SLC5 domain is competent to mediate uptake of histidine from the extracellular environment in *P. fluorescens*(*19*) and *P. putida*(*20*), and that a functional copy of CbrA is required for growth on histidine as a sole carbon/nitrogen source(19, 21). Furthermore, Wirtz *et al*. demonstrated that histidine transport through the SLC5 domain is dependent on the proton motive force(20), which is consistent with other SLC family transporters that operate through a sodium-independent transport mechanism(22–24). Although previous evidence demonstrates that the SLC5 domain of CbrA forms a functional histidine transporter(19, 20), a structural and mechanistic understanding of this histidine transport process remains unclear.

In addition to the N-terminal SLC5 transporter domain, CbrA also contains domains that extend into the cytoplasm (**Figure 1A**). Immediately following the SLC5 domain is a distinctive SLC and two-component system associated component (STAC) domain. A previous crystal structure of an isolated STAC domain from *Archaeoglobus fulgidus* revealed a small four-helical bundle(25). While the precise role of STAC domains remain unclear, their presence in various transceptor-like proteins suggests they may serve as regulatory linkers, coupling conformational changes between the membrane-embedded SLC5 domain and downstream functional regions of transceptors. Located C-terminally to the STAC domain are regions of CbrA that are classically found in histidine kinases(26), including Per-Arnt-Sim (PAS), Dimerization and Histidine phosphotransfer (DHp), and catalytic (CA) domains that extend away from the membrane (**Figure 1A**). Thermal denaturation based binding studies have previously suggested that the PAS domain in CbrA is capable of binding histidine(26), indicating that transport of histidine through the SLC5 domain may serve as a trigger for further ligand binding and/or conformational changes to activate the cytosolic histidine kinase domains. Such activation is thought to trigger phosphorylation of a conserved histidine in the DhP domain, which would then serve as a docking point for phosphotransfer to a conserved aspartate in CbrB (**Figure 1A**)(5).

Despite extensive biochemical and microbiological characterization of the CbrA/B system in various *Pseudomonas* species, the signal that triggers CbrA autophosphorylation remains elusive. It is currently unclear whether activation is driven by substrate binding and/or transport through the SLC5 domain, ligand binding to the intracellular PAS domain, coordinated conformational coupling between all domains of the protein, or some combination of all these factors(5). Progress toward mechanistic clarity on this process has largely been hampered by a lack of structural information. Beyond a crystal structure of an isolated STAC domain from an unrelated archaeal protein(25), no high-resolution structures of CbrA or its isolated domains have been reported. As a result, the molecular basis of substrate recognition, conformational communication between membrane and cytosolic domains, and the integration of these events into global regulatory outputs remain poorly understood.

Here, we address this knowledge gap by determining a cryo-electron microscopy (cryo-EM) structure of CbrA, which reveals the overall architecture of the SLC5-STAC domains, and delineates a clearly defined and conserved histidine binding pocket within the SLC5 transporter domain. Unexpectedly, a small peptide (CbrX) that is encoded immediately upstream of the transceptor remained stably bound throughout purification and cryo-EM imaging. Together with extensive molecular dynamics simulations, our studies reveal the molecular determinants of histidine binding to the SLC5 domain, identify a protonation dependent conformational switch that likely drives membrane transport, and provide a structural framework to begin understanding how membrane transport and histidine kinase signaling activities are coupled in this central regulator of *Pseudomonas* metabolism and biology.

## Results

### Expression, Purification, and cryo-EM reconstruction of CbrA

Based on previously available biochemical and microbiological data, we began our studies using the CbrA system from *P. putida* KT2440(20, 21). CbrA is highly conserved among common *Pseudomonas* species, with the protein from *P. putida* displaying 82% sequence identity to *P. aeruginosa* PAO1, and ~92% sequence identity to *P. fluorescens* (**Suppl. Figure S1**). CbrA has an overall molecular weight of 109 kDa, and in all genomes analyzed is overlapped in an operon with a small (~58 amino acid) upstream peptide called CbrX (**Figure 1B**). Previous analysis in *P. putida* has demonstrated that CbrA is translationally coupled to CbrX, with transcription of both genes being driven from a single upstream P_cbrA_ promoter(21). AlphaFold predictions suggest that CbrX adopts an alpha-helical hairpin configuration with two alpha-helices separated by a short loop (**Figure 1C**). The hydrophobic amino acid composition of the two CbrX alpha-helices suggests that the peptide may be anchored within the lipid membrane. However, the cellular localization and overall functional role of CbrX beyond transcriptional coupling to CbrA has yet to be determined.

To facilitate expression and purification of CbrA for structural studies, we constructed a pETDuet-1 expression vector containing full-length CbrX and CbrA genes from *P. putida* each driven by individual T7 promoters (**Suppl. Figure S2A**). CbrA was tagged with an 8x-histidine tag at the C-terminus for purification purposes, and CbrX was left untagged. Following heterologous expression of CbrX and CbrA together from this single plasmid in C41(DE3) *E. coli,* isolated membrane fractions were solubilized in lauryl maltose neopentyl glycol (LMNG) detergent and CbrA was purified by two-step Co^3+^-Talon affinity and size-exclusion chromatography. The resultant size-exclusion chromatogram and SDS-PAGE of peak fractions revealed a relatively monodisperse preparation of full-length CbrA at the expected molecular weight of ~109kDa (**Suppl. Figure S2B-D**). Based on calibrated molecular weight standards, the elution volume of CbrA on a Superdex 200 Increase 10/300 GL size-exclusion column (~11.8 mL) was most consistent with a monomer of CbrA plus the associated mass of an LMNG micelle.

As an initial screening step, the detergent solubilized preparations of purified full-length CbrA were plunge-frozen on cryo-EM grids and imaged on a Talos Arctica 200 keV electron microscope. The resultant micrographs displayed an even distribution of seemingly small particles in thin ice (**Suppl. Figure S2E**). 2D averages calculated from particles extracted from these micrographs show a small membrane protein embedded within a detergent micelle, with a small protrusion from the detergent micelle corresponding to the intracellular STAC domain (**Suppl. Figure S2F**). From the 2D averages alone it is readily apparent that detergent solubilized full-length CbrA is purified in monomeric form consistent with the elution volume observed with size-exclusion chromatography, and also that the cytosolic PAS-DHp-CA domains are completely disordered and not visible in the averages (**Suppl. Figure S2F**). Based on this preliminary screening dataset we constructed a truncated version of CbrA in which the flexible and disordered PAS-DHp-CA domains were removed, leaving an expression plasmid with only CbrX and the SLC5-STAC domains of CbrA which encompass a calculated protein mass of ~66 kDa. Purification of this truncated construct resulted in significantly improved overall protein yield and monodispersity as assessed by size-exclusion chromatography (**Suppl. Figure S3A-C**).

For high-resolution cryo-EM studies we prepared plunge frozen grids of the truncated SLC5-STAC CbrXA construct which were imaged at 300 keV on a Titan Krios G4i. The resultant dataset revealed particle distribution and calculated 2D averages (**Suppl. Figure S3D&E**) that were highly similar to those obtained with the full-length CbrA construct, further indicating that the PAS-DHp-CA domains are disordered in the full-length construct. Subsequent 3D classification and reconstruction (**Suppl. Figure S4**) revealed a predominant single conformation of the CbrA SLC5-STAC domains which was reconstructed to a final overall resolution of 1.9Å (**Figure 2A**, **Suppl. Figure S5A-D, and Suppl. Table S1**). At such resolution the individual rotamer states of most residues are easily discerned (**Suppl. Figure S5E**), and holes are observed in the coulomb potential map for most aromatic sidechains (**Suppl. Figure S5F**). The high resolution of the final map greatly facilitated refinement of an atomic model, and also revealed several well-ordered water molecules throughout different regions of the SLC5-STAC domain (**Suppl. Figure S5G**).

**Figure 2.**
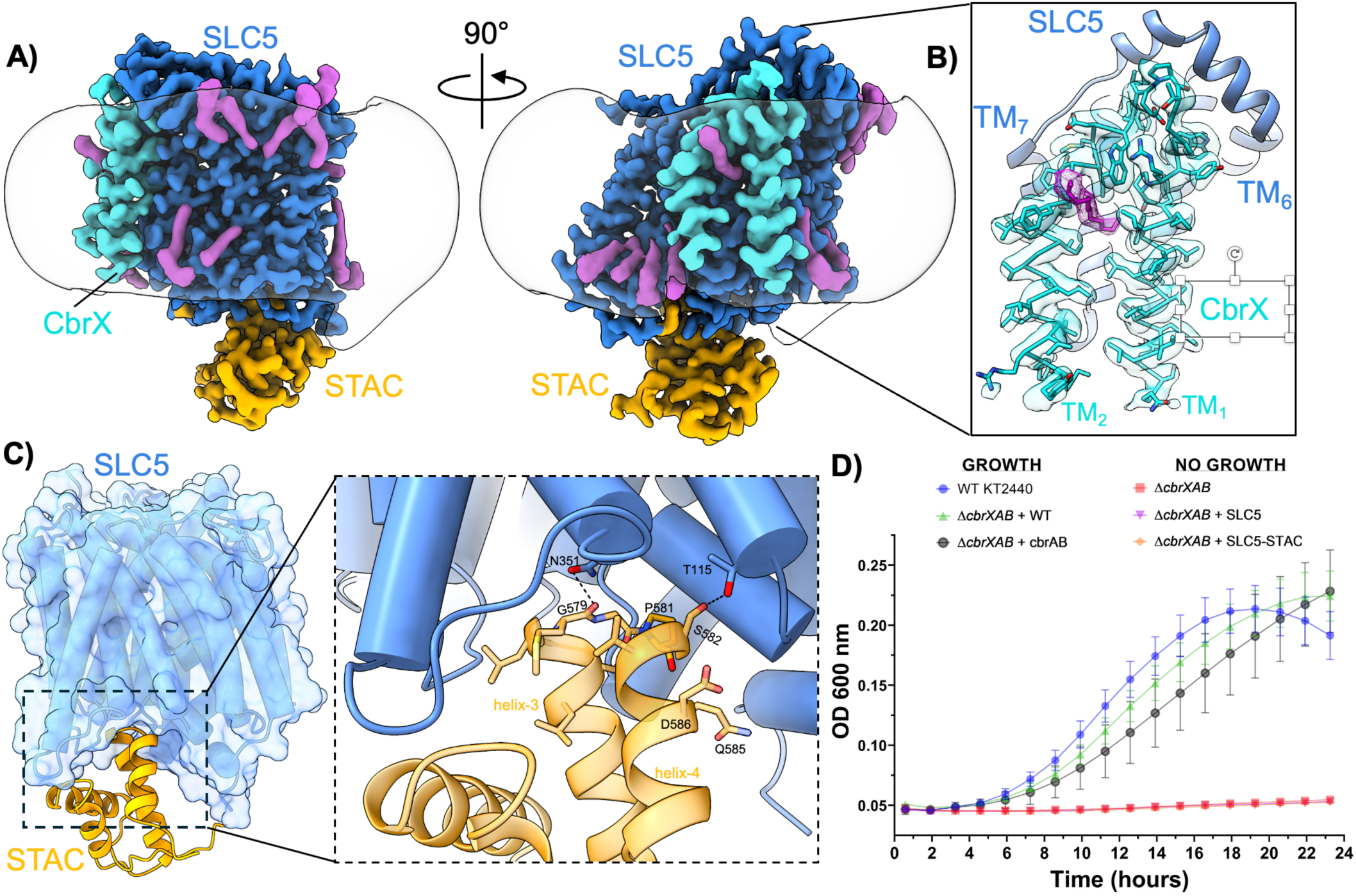
Structural analysis of CbrA SLC5-STAC construct. **A)** Rotated views of the high resolution cryo-EM map of the CbrA SLC5-STAC domains. The SLC5 domain is colored blue, STAC domain is colored orange, and CbrX is shown in cyan. Copurified lipid/detergent moieties are shown in magenta. The boundary of the detergent micelle is shown as a transparent white surface. **B)** Zoomed in view of co-purified CbrX demonstrating clear sidechain density throughout the peptide. A lipid like density (magenta) inserts between the two helices of CbrX. **C)** Diagram showing the interaction between SLC5 and STAC domains in CbrA. Coloring scheme is the same as in **A**. Dashed box inset provides a zoomed in view of the specific interactions between SLC5 and STAC domains. Hydrogen bonds are shown as dashed black lines. **D)** Growth curves of *P. putida* in M9 minimal media with histidine as a sole carbon and nitrogen source. CbrX is not required for growth on histidine. The SLC5 or SLC5-STAC constructs which lack intracellular histidine kinase domains of CbrA are not sufficient to rescue growth on histidine. Data points represent the average of three (N=3) biological replicates that were each collected in technical triplicate. Error bars represent standard error of the mean (SEM) among biological replicates.

### Structure of the CbrXA SLC5-STAC domains

The high-resolution cryo-EM map of CbrA SLC5-STAC domains revealed all 13 TM helices of CbrA, with several copurified lipid/detergent like densities also visible around the periphery of the transmembrane region (**Figure 2A**). Most surprisingly, clear density for the short CbrX peptide was also present in the final reconstructed cryo-EM map (**Figure 2A&B**), indicating that CbrX forms a stable interaction with CbrA even after solubilization from membranes with detergent. In the reconstructed map CbrX adopts an alpha-helical hairpin configuration with both the amino and carboxy termini facing the intracellular region, and a short loop that connects both helices facing the periplasmic space. CbrX was positioned with both helices packed against TM_3_ and TM_8_ of the CbrA SLC5 domain, and underneath a structured alpha helical loop in the periplasmic space that connects CbrA TM_6/7_ (**Figure 2A&B**). Within the cryo-EM map a thin acyl chain-like density likely corresponding to a co-purified lipid or detergent molecule is observed inserted between the two CbrX TM helices (**Figure 2B**). As elaborated upon in the discussion, the position of this lipid-like density suggests a possible explanation for purification of CbrA in a monomeric form, rather than as a dimer which is typical of most histidine kinase family proteins.

The STAC domain was also directly visible in the final cryo-EM map and adopted a four-helix bundle similar to the previously determined crystal structure of an isolated STAC domain from *A. fulgidus*(*25*). The STAC domain is connected to the SLC5 domain through a 30-residue linker extending from TM_13_ in the SLC5 domain. The N-terminal region of this linker (residues 497-507) is ordered as an alpha-helix that runs parallel to the membrane plane, with the remainder (residues 509-517) disordered and not visible in the map. The STAC domain is positioned beneath the membrane plane with the tips of STAC domain helices 3 and 4 inserted into a pocket formed at the base of the SLC5 transporter domain (**Figure 2C**). Most of the interactions between the SLC5 and STAC domains are hydrophobic in nature, with hydrogen bonds also observed between N531 and the backbone carbonyl of G579, and the sidechain hydroxyl of T115 and S582 (**Figure 2C**). Together, these interactions anchor the STAC domain against the cytoplasmic face of the SLC5 transporter domain in a defined orientation, suggesting that STAC serves as a structurally integrated appendage rather than a flexibly tethered accessory. This positioning suggests that the STAC domain may sense conformational changes within the SLC5 transporter core and potentially relay them to the downstream histidine kinase domains.

To assess the functional significance of CbrX and different domains of CbrA we constructed a markerless knockout of the entire CbrXAB operon in *P. putida* KT2440 using allelic exchange(27). Consistent with previous studies, growth of wild-type and *ΔcbrXAB* strains in defined minimal media revealed that *cbrXAB* is essential for growth when L-histidine is supplied as the sole carbon and nitrogen source(19, 21) (**Figure 2D**). Complementation of the knockout strain was achieved by mini-Tn7 based insertion of a single copy of the wild-type *cbrXAB* operon with native promoters at the *att*Tn7 site downstream of the *glmS* gene(28), which complemented growth on histidine as a sole carbon and nitrogen source back to levels seen with wild-type *P. putida* (**Figure 2D**). When the *ΔcbrXAB* knockout strain was complemented with a *cbrAB* operon lacking the gene encoding the CbrX peptide, growth was also restored to levels comparable to the wild-type strain (**Figure 2D**). This result demonstrates that while CbrX is translationally coupled to CbrA(21), production of the CbrX peptide is not required to produce a functional copy of CbrA and complement growth on L-histidine. Similarly, while previous studies have demonstrated that the SLC5 domain of CbrA alone can mediate histidine uptake in *P. putida*(*20*), the functional significance of such uptake remained unclear. When we complemented the *ΔcbrXAB* strain with a *cbrXAB* operon that contained only the SLC5 or SLC5-STAC domains of CbrA, no growth on histidine as a sole carbon/nitrogen source was observed (**Figure 2D**). These results indicate that any histidine transport capacity of the SLC5 or SLC5-STAC domains is insufficient for growth of *P. putida* on histidine and that a full-length CbrA protein complete with intracellular histidine kinase domains is required for complementation, which corroborates findings in *Pseudomonas fluorescens* SBW25(19).

### Histidine binding in the SLC5 domain

Previous studies have demonstrated that the CbrA SLC5 domain can bind histidine and mediate proton gradient driven uptake of the amino acid from the extracellular environment(20). In our cryo-EM map the TM helices of the SLC5 domain are arranged in a classic LeuT-type membrane transporter fold (**Figure 3A&B**)(29). TM_2,3,7,8_ and TM_4,5,9,10_ form “bundle” and “hash” subdomains, with TM_6,11_ forming gating helices on the periphery of the transporter core(30). The TM helices are arranged with TM_3-6_ and TM_8-11_ in an inverted repeat architecture, with helical breaks in TM_2_ and TM_7_ forming a central transport substrate pocket analogous to that seen in other LeuT type transporters(29–31). The cryo-EM map revealed a distinct density within these helical break regions that is not attributable to CbrXA, and is consistent with the size and shape of a free histidine molecule (**Figure 3C&D and Suppl. Figure S6A&B**). Analysis of the cryo-EM map at different threshold levels reveals that the ligand density is significantly weaker than the surrounding protein density, suggesting that the copurified ligand is present at reduced occupancy relative to the protein and/or that its binding within the pocket is weakly constrained by surrounding residues (**Suppl. Figure S6C**). Despite this potential reduced occupancy or weak binding, based on the shape of the observed density, it’s location in the canonical LeuT-type transport pocket, and previous reports of histidine transport by CbrA(19, 20), we have modeled this extra density as a co-purified free histidine molecule (**Figure 3C and Suppl. Figure S6**).

**Figure 3.**
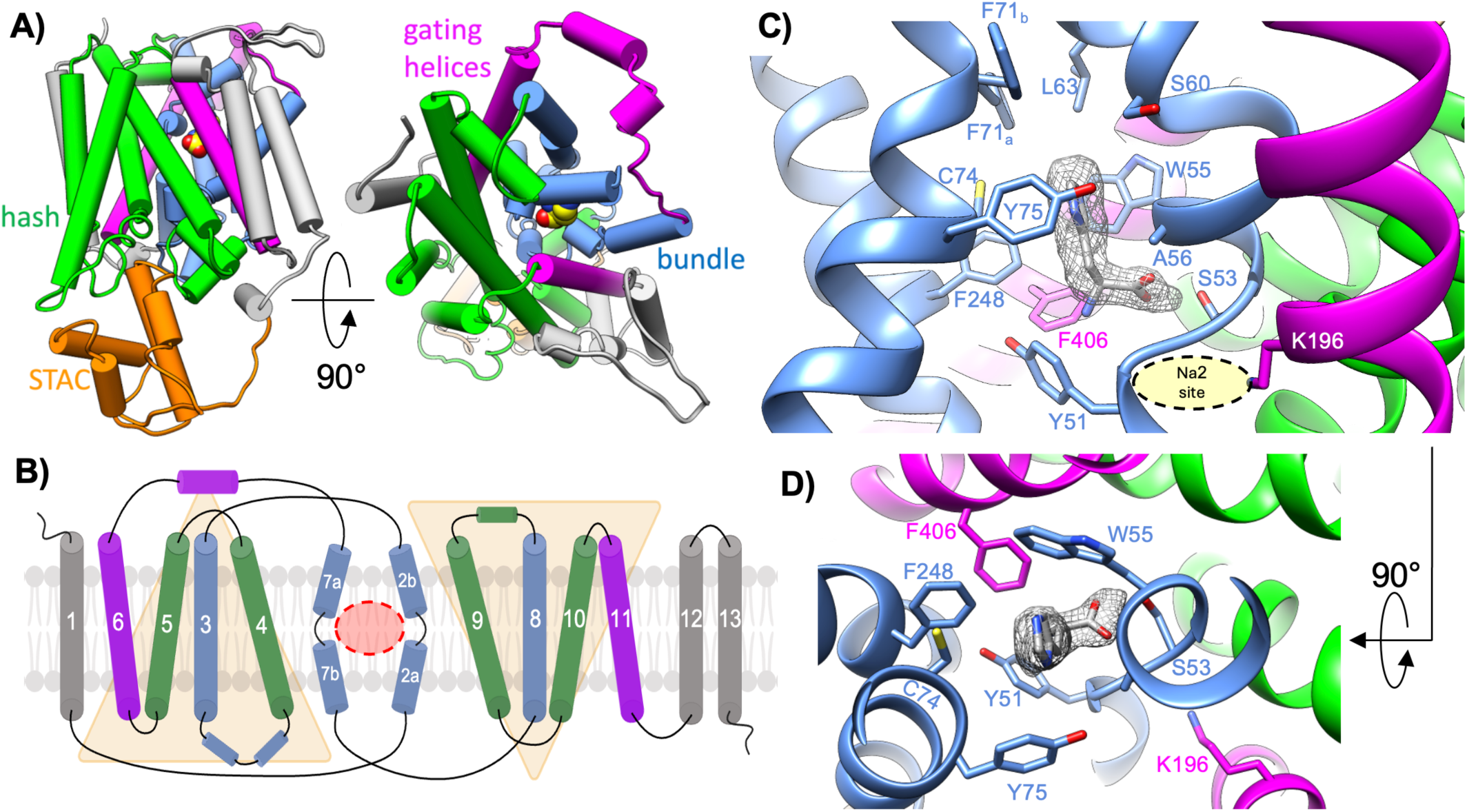
Histidine binding pocket within the SLC5 domain. **A)** Overall topology of the CbrA SLC5 domain colored according to conventional LeuT type transporter folds. TM helices corresponding to the hash subdomain are colored green, the bundle subdomain colored blue, and the gating helices colored magenta. Extra TM helices outside the conventional LeuT topology are colored grey. The STAC domain is shown in orange beneath the SLC5 domain. **B)** Topology diagram of the SLC5 TM helices colored the same as in **A**. TM_3-6_ and TM_8-11_ form two inverted repeats highlighted with transparent yellow triangles. Helical break regions are identified in TM_2_ and TM_7_, which form the central ligand binding pocket outlined in a dashed red circle. **C)** View of the central transport pocket in the SLC5 domain showing density for a copurified molecule of the approximate size and shape of histidine. Individual TM helices are colored as in pane **B**. The Na2 site characteristic of LeuT type transporters is occupied by the sidechain of K196. **D)** Rotated view of panel **C** showing the histidine binding pocket as viewed from the periplasmic space.

The co-purified histidine captured within the center of the CbrA SLC5 domain is oriented with the imidazole ring moiety pointing toward the periplasmic space (**Figure 3C&D, Suppl. Figure S6**). The substrate binding pocket is lined by several hydrophobic sidechains, including A56, W55, Y51, Y75, F248, and F406 of CbrA (**Figure 3C&D, Suppl. Figure S6A&B**). Just above the bound histidine we observed two alternate rotamer states of the F71 sidechain, one of which partially seals the substrate binding pocket from the periplasmic space (**Suppl. Figure S6D**). The carboxylate moiety of the bound histidine interacts closely with S53 in the helical break region of CbrA TM_2_ (**Figure 3C, Suppl. Figure S6A**). On the opposite side of this helical break is the canonical Na2 site found in various sodium coupled LeuT type transporters(30). Consistent with previous reports of histidine transport by CbrA being independent of a sodium gradient and dependent on the proton motive force(20), the Na2 site in CbrA is occupied by the sidechain of K196 (**Figure 3C&D**). The pKa of the K196 sidechain amino group as calculated with PROPKA(32) is 7.36, consistent with the lower pKa seen with similarly placed lysine sidechains in other proton coupled LeuT type transporters(22, 24). The low pKa of the K196 sidechain suggests that this residue may act as a titratable residue to bias the conformation of the helical break region in TM_2_ similar to Na^+^ binding in sodium dependent transporters(30), thus modulating SLC5 domain dynamics for histidine transport.

### Water permeation in the CbrA SLC5 domain

In addition to greatly facilitating refinement of an atomic model, the high resolution obtained from 3D reconstruction of the CbrA SLC5-STAC domains revealed many water molecules directly visible in the cryo-EM map (**Figure 4A and Suppl. Figure S5G**). Several well-ordered water molecules are observed within the vicinity of the bound histidine, and also within the Na2 site near the helical break in TM_2_ that is occupied by the K196 sidechain (**Figure 4B**). Many water molecules are also observed in the cytoplasmic vestibule where the STAC domain interacts with the hash subdomain of the SLC5 transporter (**Figure 4C**). The presence of these well-ordered water molecules in the cryo-EM map demonstrates that a significant proportion the transport pathway through the SLC5 transporter core remains solvated following solubilization in detergent.

**Figure 4.**
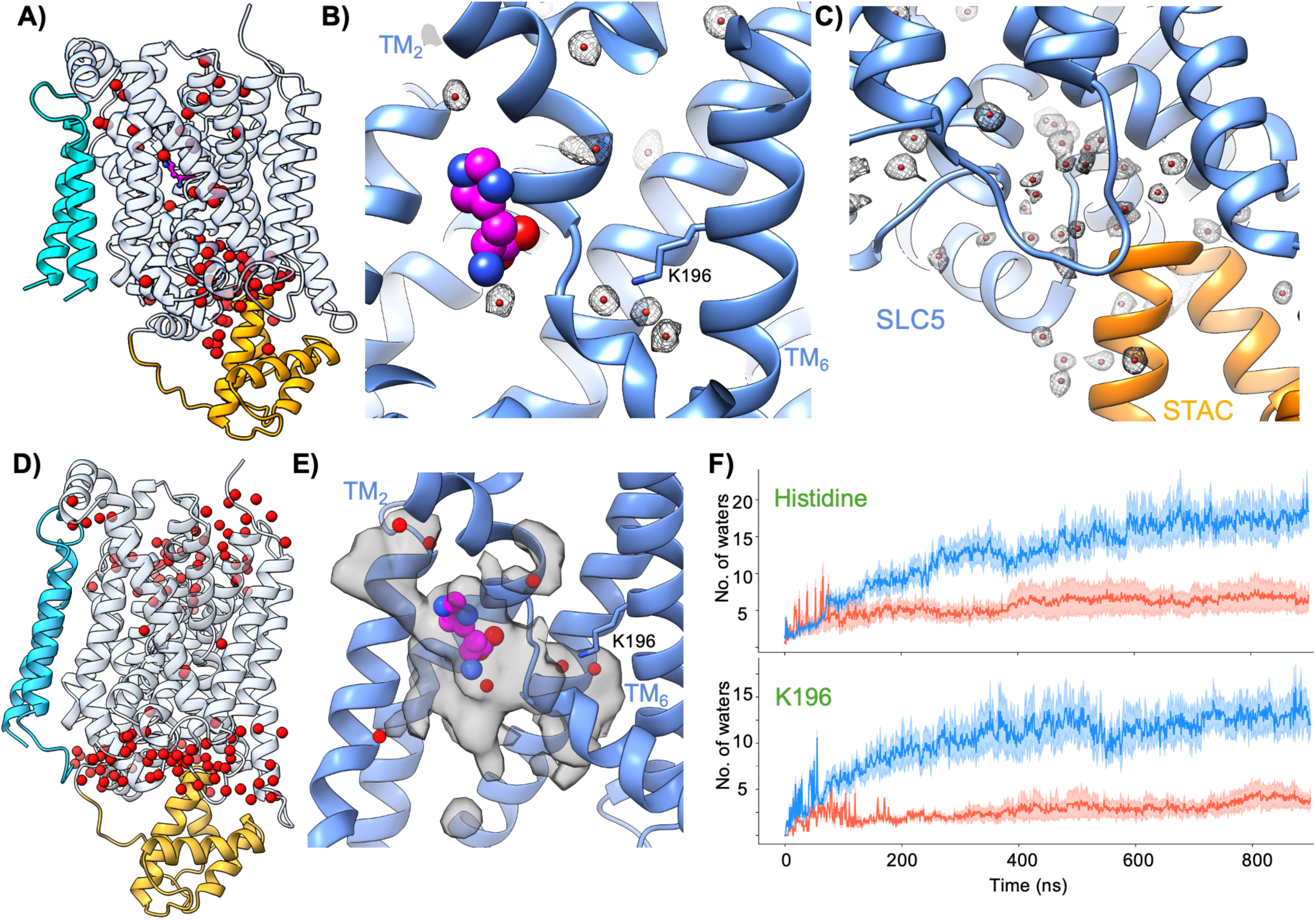
Water permeation analysis. **A)** Waters observed in the cryo-EM map of the CbrXA SLC5-STAC domains. Waters are shown as red spheres. CbrX is colored cyan, the CbrA SLC5 domain is shown in transparent light blue, and the STAC domain is shown in orange. **B)** Close-up view of the cryo-EM resolved waters positioned near K196 in TM_6_, and the periphery of the histidine binding pocket. Waters are shown as red spheres with the corresponding coulomb potential map shown in grey mesh. **C)** View of waters (red spheres) resolved in the cryo-EM map (grey mesh) in the cleft beneath the SLC5 domain (blue) that is occupied by the STAC domain (orange). **D)** Single snapshot from one of the MD simulation trajectories with the CbrXA SLC5-STAC domains and waters within 3 Å of the protein. Coloring scheme is the same as in panel **A**. **E)** Close-up view of HIS binding pocket, TM_6_ (K196) and TM_2_ (break region) with the volume map of water occupancy from MD simulations shown in black transparent solid. The volume map is overlayed on the cryo-EM structure with waters shown in red. The occupancy map of dynamic waters (MD) completely confine all the static waters observed in the cryo-EM map. **F)** Time evolution of average water occupancy within 1 nm of the CA atom of histidine (top panel) and K196 residue across protonated (in red) and deprotonated (in blue) MD simulations. The shaded region shows standard error of mean across all five replicates.

While the cryo-EM map provides sufficient resolution to identify bound water molecules, the inherently static nature of the experiment does not inform on the dynamic exchange of water into and out of distinct regions of the transporter. To gain insight into the conformational dynamics of CbrXA and the permeation of water within the SLC5 transporter core, we performed all-atom molecular dynamics (MD) simulations of the CbrA SLC5–STAC domains embedded in a lipid bilayer. A visual representation of the simulation box with all the components are shown in supplemental data (**Suppl. Fig. S7A**). Due to the near-neutral pKa of the conserved K196 that is central to the Na2 site, we decided to carry out two separate sets of MD simulations with the K196 side-chain held fixed in either protonated or deprotonated state, treating protonation as a controlled perturbation rather than a dynamically sampled variable. For each of these two systems, henceforth denoted as “K196 protonated” and “K196 deprotonated”, we simulated five independent replicates totaling approximately 5 μs of aggregate simulation time for each. Importantly, the water molecules observed in the cryo-EM map were not included in the initial setup of the simulation systems, and water was instead restricted to the bulk aqueous regions outside the lipid membranes while setting up the molecular dynamics simulations. Hence, the water molecules that permeated the CbrA transporter core in the MD simulations originated from the external solvent.

Analysis of the MD trajectories revealed that water molecules consistently permeated into the core of the CbrA SLC5 domain and occupied positions that closely correspond to those observed in the cryo-EM structure. The permeated water molecules formed hydrogen-bonded networks extending from the bulk solvent towards the helical break in TM_2_ around the histidine binding pocket and K196 residue (**Suppl. Fig. S7B-D**). A representative MD snapshot (**Figure 4D**) of water molecules inside CbrA was extracted once a steady state of water influx had been attained. In this static picture, we observed strong correlation between the positions of permeated waters in simulations with that of the stably bound waters in the experimental cryo-EM structure. Subsequently, utilizing the MD trajectories we constructed a density map of permeated waters in the SLC5 domain (**Figure 4E**). This dynamic representation revealed a strong spatial overlap between the probability density of waters seen in MD with that of cryo-EM resolved waters. Furthermore, water molecules consistently infiltrated the Na2 site region surrounding the conserved K196 residue and the histidine-binding pocket in MD simulations, albeit with different frequencies in protonated and deprotonated systems (**Suppl. Fig. S7C&D**). The time evolution of water permeation in CbrA (**Figure 4F**) showed that the average number of waters within 1 nm of the C*α* atom of K196 (top panel) or histidine substrate (bottom panel) is typically 3-5 times higher in K196 deprotonated simulations compared to its protonated counterpart. This pattern is prominent across all replicates once a steady state of water influx has been achieved, typically around the 400 ns timescale. The preferential hydration in deprotonated K196 systems indicates a relationship between the protonation state of the K196 and water permeation through the transporter core, hinting at a potential impact in the histidine transport mechanism.

Overall, the strong correspondence between water molecules identified experimentally by cryo-EM and those sampled in the dynamic MD trajectories (**Figure 4-E**) indicates that hydration within the CbrA SLC5 transporter represents an intrinsic and dynamically accessible feature of the protein rather than an artifact of experimental preparation. Water permeation near the Na2 site, captured in both cryo-EM and MD simulations, suggests that the local environment surrounding K196 could support plausible pathways for solvent-mediated proton access. Such proton access could plausibly lead to biasing of the K196 protonation state, leading to altered conformational states of the transporter to support histidine transport.

### K196 (de)protonation and its effect on CbrA conformation space

Given the conserved placement of K196 within the Na2 site of the CbrA SLC5 domain and the intriguing observation of water permeation near this residue in CbrA, we decided to examine whether the protonation state of K196 would also influence the conformational landscape of the SLC5-STAC domains and have an effect on histidine transport, as has been observed for similarly placed lysine residues in other proton-coupled SLC transporters(22–24). The previously mentioned MD simulation trajectories of the CbrXA SLC5–STAC domains in either protonated or deprotonated K196 state, were analyzed to understand the histidine motion and relevant conformational changes. First, to assess the stability of binding and probable transport of histidine in CbrA, we estimated the root mean square deviation (RMSD) of histidine coordinates in both protonated and deprotonated MD trajectories (**Figure 5A**). The average RMSD across five replicate simulations revealed noticeable differences in histidine motion between the K196 protonated and deprotonated systems, with the deprotonated systems showing a steady increase in the RMSD post 600 ns compared to the protonated systems. Next, we computed the average RMSD of the TM_2_ break region heavy atoms (**Figure 5B**) to understand the motion of CbrA in each of the two systems. Consistent with histidine deviation, we noticed that the deprotonated ensembles have a 2-fold higher RMSD in the break region compared to the protonated ensembles. Interestingly, both histidine and break region RMSDs in deprotonated simulations typically start to show this upward trend around 500-600 ns, after the steady-state of water permeation is achieved (~ 400 ns), which may point to a subtle correlation between the K196 protonation dependent hydration with that of the conformational dynamics of the transporter.

**Figure 5.**
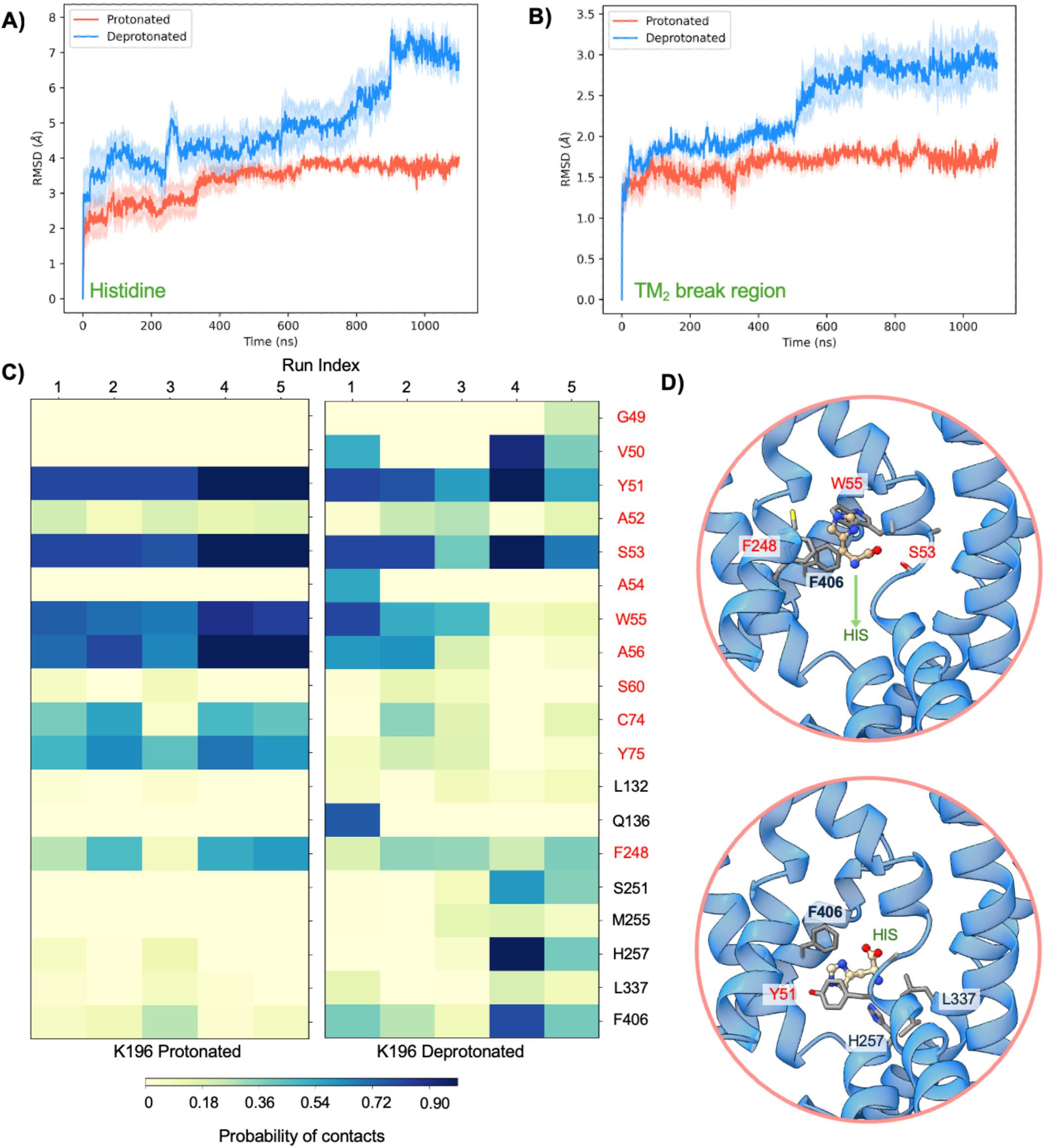
RMSD and residue contact analysis from MD simulations. Average root mean square deviations (RMSD) of histidine **(A)** and TM_2_ break region **(B)** across five replicates of protonated (blue) and deprotonated (red) simulations with shaded areas showing the standard error across 5 replicate simulations. **C)** Heatmap of intermolecular interactions between histidine and CbrA residues across five replicates of protonated and deprotonated simulations. The color scheme represents the percentage of snapshots in a replicate that have contact between histidine and the specific amino acid. Amino acids labeled in black primarily form contacts in deprotonated CbrA simulations. **D)** Representative snapshots from MD trajectories highlighting different histidine-CbrA interactions near the binding pocket (top panel) and after the downward transport (bottom panel).

Examining the intermolecular interactions between the bound histidine molecule and CbrA from the MD trajectories highlights significant residues that contact histidine at different timepoints of the trajectories. Histidine-CbrA interactions were quantified across five replicates of protonated and deprotonated K196 simulations (**Figure 5C**). The set of *significant residues* is determined by a threshold that the residue and histidine contact must be present in at least 10% of the frames in either of the two systems, and *contact* is defined when the histidine:CbrA heavy atom distances are less than 3.5 Å (33, 34). The contact maps of the protonated simulations show high probability of histidine interactions with Y51, S53, W55, A56, C74 and Y75 residues (**Figure 5C, and 5D upper panel**). However, this pattern is significantly shifted in deprotonated simulations, where unique interactions with F248, S251, H257, and F406 were observed in our MD trajectories that were not seen in protonated counterparts (**Figure 5C, and 5D lower panel**). It should be noted that these new contacts in the deprotonated system are only possible after substantial displacement of histidine and considerable motion in the break region of CbrA TM_2_.

Histidine transport mediated by CbrA is a complex process that likely requires intricate, long timescale motion in the protein to attain favorable conformations for transport. To examine the slow motions in the conformational space of the histidine bound CbrA, we carried out combined time-lagged independent component analyses (tICA) with all the simulation data. In this dimensionality reduction technique, the slow conformational motions are resolved into independent components (collective modes) enabling us to study progress along these slower modes(33, 34). We trained 2-dimensional tICA models using the combined dataset of protonated and deprotonated simulation features, one each with histidine coordinates and aligned break region heavy atom coordinates respectively. The tICA models are trained with a combined dataset (protonated and deprotonated) to ensure consistency of the definitions of independent components across both systems. We compared the free energy distributions in the tICA space between protonated and deprotonated systems with histidine positions as features towards the tICA model (**Figure 6A**). The protonated ensembles have two energetically stable, localized, free energy minima (a) and (b), where the backbone of S53 interacts with histidine by forming H-bonds with the carboxyl moiety of the ligand. In each of these energetic basins the histidine molecule is oriented in different directions (**Figure 6C**). On the other hand, deprotonated ensembles have three relatively spread-out shallow basins. In general, the free energy distribution of the deprotonated system showed a relatively high spread of conformational sampling. Interestingly, one of the stable basins (d) captures histidine in a slightly lower pocket along the channel enabling an H-bond between the ligand amino group and the S53 backbone carbonyl.

**Figure 6.**
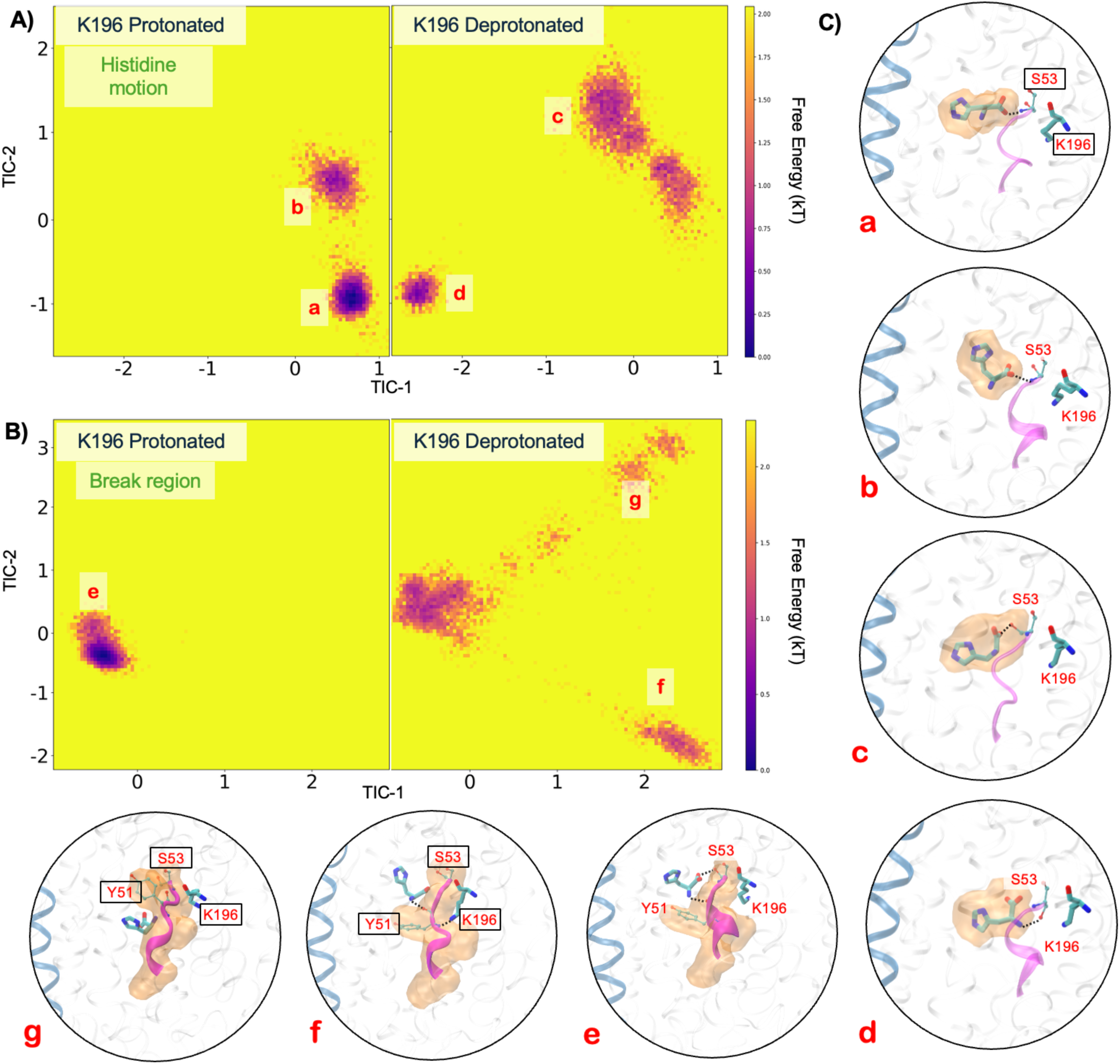
Time-lagged independent component analysis. Free energy distributions along tIC-1 and tIC-2 with **(A)** histidine coordinates as a feature, and (**B**) heavy atoms of the break region as a feature for tICA training in protonated and deprotonated systems. **C)** Representative conformations and important residues captured in highlighted energy basins are shown. Red lowercase letters correspond to the energy basins highlighted in panel **A** and **B**. The TM2 break region is shown in purple in all snapshots with Histidine and K196 shown in licorice. Orange transparent surface corresponds to the average position of histidine (basin a-d) or the TM_2_ break region (basin e-f) across 5 replicate MD simulations.

From the second tICA model, trained on the heavy atom coordinates of the break region residues, we observed one remarkably stable free energy minima (e) in the protonated ensembles suggesting no significant motion in the break region. In fact, we noticed the break region forming a stable helical turn in >50% of the protonated ensembles. Also, in the protonated system the Y51 sidechain forms a barrier at the base of the histidine binding pocket, preventing diffusion of the ligand to the intracellular space (**Figure 6E, panel e**). Interestingly, the deprotonated system demonstrated enhanced flexibility of the break region with three distinct basins and internal conversions. While we observed a broad, sufficiently stable basin in similar free energy space as the one in the protonated system, the other two shallower basins appear at the two extremes of tIC-1 revealing unique ensembles of conformations. The break region attains a flexible loop conformation in both (f) and (g) basins. Remarkably, the ensemble (g) showed a considerable conformational shift in the break region, in particular the sidechain of Y51, which shifted away from its native gating location. This resulted in substantial downward transport of the HIS molecule toward the intracellular space, correlating to an RMSD of 10 Å in the histidine molecules in this basin compared to basin (e) or (f). These findings suggest that the protonation state of K196 can modulate the flexibility of the break region and subsequent orientation of Y51 as one component of enabling conformational transitions to mediate histidine transport through CbrA.

Together, our cryo-EM structure and molecular dynamics simulations reveal that the CbrA SLC5 domain retains hallmark features of proton coupled amino acid transporters(22, 24). These features include a conserved Na2 site that is occupied by a lysine residue with a near neutral pKa that likely facilitates amino acid transport by biasing transporter conformations through a (de)protonation mechanism.

## Discussion

The ability of *Pseudomonadaceae* to thrive across diverse ecological niches relies on their capacity to sense and metabolize fluctuating nutrient sources. Central to this adaptability is CbrA, an unusual fusion protein that combines an SLC5-family membrane transporter with a cytosolic histidine kinase, enabling coordinated regulation of carbon and nitrogen metabolism(5). Although CbrA is essential for the utilization of substrates such as histidine, how membrane transport events are mechanistically coupled to downstream signaling and metabolic control has remained poorly understood(19, 20). Here, we addressed this gap by determining a high-resolution cryo-EM structure of the CbrA SLC5–STAC domains, revealing the architecture of the membrane transporter region and a conserved histidine-binding pocket. Integrated with all-atom molecular dynamics simulations, these data provide a framework to begin understanding how ligand recognition and transporter conformational dynamics may be communicated across domains to allow CbrA to function as a global metabolic regulator in *Pseudomonadaceae*.

CbrA is translationally coupled to the short peptide CbrX(21), and our cryo-EM structure reveals that CbrX is a membrane integrated peptide that forms a stable interaction with the CbrA SLC5 transporter domain (**Figure 2A&B**). Although histidine kinases classically function as dimers(26), heterologously expressed CbrXA purified in detergent was predominantly monomeric (**Suppl. Figure S2&S3**). In contrast, AlphaFold models predict a dimeric configuration of CbrA that is consistent with canonical histidine kinase architectures (**Suppl. Figure S8A&B**), in which two CbrX peptides appear to mediate contacts between opposing CbrA SLC5 domains (**Suppl. Figure S8C**). We suspect that detergent solubilization weakens interactions between CbrX peptides at the dimer interface between opposing CbrA monomers, leading to the monomeric preparations of CbrXA obtained here. This idea is supported by the observation of acyl-chain like density interdigitating between the TM helices of CbrX (**Figure 2B**). Together, these observations suggest that a native membrane environment and specific lipid interactions may be required to stabilize the functional oligomeric state of CbrXA. However, it is important to note that *P. putida* growth assays demonstrate that CbrX is not essential for CbrA-dependent growth on histidine as a sole carbon and nitrogen source (**Figure 2D**). Thus, while CbrX co-purifies with CbrA and may contribute to dimer stabilization, its precise mechanistic role in CbrA signaling remains unresolved. Future biophysical assays with elaborate membrane reconstitution methods and/or *in vivo* assays will be required to investigate the oligomeric state of CbrXA in a (near)native membrane context and assess the mechanistic role of CbrX.

The high-resolution structure of the CbrA SLC5–STAC domains presented here provides a framework for understanding how histidine sensing and transport are mechanistically encoded within the transmembrane region of CbrA. Our cryo-EM structure revealed that the SLC5 domain of CbrA forms a classical LeuT type transporter topology with a histidine binding pocket that is analogous to the transport pocket identified in other SLC family amino acid transporters (**Figure 3A-C**) (22, 24). The CbrA K196 sidechain is pointed into the Na2 site characteristic of other proton coupled SLC transporters (**Figure 3C&D**), with a calculated pKa that is consistent with this residue likely facilitating protonation dependent conformational transitions of a key helical break in TM_2_ to mediate histidine transport(22, 24). Molecular dynamics simulations with protonated and deprotonated K196 supported this hypothesis as significant deviations in the helical break of TM_2_ were observed only when K196 was deprotonated (**Figure 5B**). This deviation in the TM_2_ helical break aided in a relatively higher movement and a downward motion of the histidine substrate towards the cytosol. The free energy landscapes along two of the slowest conformational modes in tICA space (**Figure 6A&B**) shows that both histidine and the TM_2_ break region sample a diverse ensemble of conformations in when K196 is deprotonated compared to protonated. The break region, in its fully flexible loop structure, facilitated the gating residue (Y51) to adopt an alternate conformation that allows the histidine to transition significantly towards the cytoplasm. These free energy landscapes along slower modes provide a preliminary mechanistic understanding of a long time-scale histidine transport process. MD simulations indicate an increased structural plasticity of the break region and substrate histidine with substantially more waters permeating in deprotonated simulations. Previous studies have demonstrated that in other proton coupled amino acid transporters such as MjApcT(22), HutT(23), and GkApcT(24), a lysine pointing into the conserved Na2 site is essential for substrate transport. Analysis of mutational effects on CbrA in *P. putida* are complicated by the fact that CbrA signaling induces expression of the high-affinity histidine transporter HutT, which serves as the primary histidine importer (**Figure 1A**)(23). Thus, even partial signaling by CbrA variants may therefore be sufficient to activate HutT expression, masking potential defects in CbrA-mediated histidine transport. Furthermore, both CbrA (**Figure 2D**) and HutT (23) are essential for growth of *P. putida* on histidine as a sole carbon source, which significantly complicates development of experiments to assay CbrA histidine transport capacity in a native context. These observations highlight the need for *in vitro* assays that decouple CbrA transport activity from downstream transcriptional responses. Although extensive efforts by us and other groups to functionally reconstitute CbrA into proteoliposomes have thus far been unsuccessful (19, 20), future advances in membrane reconstitution strategies and/or modulation of lipid composition may enable direct interrogation of CbrA transport and signaling mechanisms in a controlled *in vitro* system. Such approaches will be critical for defining how specific residues and domain motions contribute to the function of this unusual transporter-kinase fusion.

Mechanistic interpretation of CbrA function is strengthened by the resolution of our cryo-EM reconstruction. Achieving sub–2Å resolution cryo-EM reconstructions (**Suppl. Figure S5**) for a ~66 kDa membrane protein assembly is rare, and enabled direct visualization of features typically inaccessible in cryo-EM studies of small membrane transporters. This level of detail allowed visualization of ordered water molecules throughout the CbrA SLC5–STAC domains, including within the histidine-binding pocket, surrounding the titratable K196 residue, and extending into a solvent-accessible vestibule beneath the SLC5 domain where the STAC domain docks (**Figure 4A-C and Suppl. Figure S5G**). The presence of structured waters in these regions suggests potential pathways for proton and solvent coordination that may facilitate conformational transitions during transport and signaling. Consistent with this interpretation, all-atom molecular dynamics simulations revealed dynamic water permeation into these same cavities (**Figure 4D-F**), supporting the notion that hydration plays a role in stabilizing intermediate conformational states of the transporter. Together, the convergence of high-resolution structural data and molecular dynamics affirms the importance of solvent-mediated interactions in shaping the functional landscape of the CbrA transmembrane region.

In our cryo-EM reconstruction the STAC domain forms an extensive and well-ordered interface beneath the SLC5 transporter core. This positioning places the STAC domain in an ideal location to sense and respond to conformational changes within the SLC5 transporter domain. Rather than serving as a passive linker, the tight packing and geometry of the STAC–SLC5 interface suggest that the STAC domain may act as a mechanical or allosteric coupling element, relaying transporter motions toward the downstream histidine kinase domains. Subtle rearrangements within the SLC5 domain that are driven by ligand binding, protonation state, or hydration changes, could plausibly be transmitted through the STAC domain to modulate kinase activity. This architecture supports a model in which CbrA operates as a true transceptor, integrating transport-derived conformational signals directly into regulatory outputs, and provides a structural basis for how membrane-localized events may control cytosolic signaling in this system.

In summary, our work provides a high-resolution structural and dynamic framework for beginning to understand how CbrA integrates membrane transport with global metabolic regulation in *Pseudomonadaceae*. By revealing the architecture of the SLC5–STAC domains, identifying a conserved histidine-binding pocket, and uncovering hydration and protonation-dependent conformational dynamics within the transporter core, this study establishes key molecular features that underlie the overall function of CbrA. The close association of the STAC domain with the SLC5 transporter further suggests a direct route for coupling transport-derived motions to histidine kinase activation, offering new insight into how transceptors may have evolved to coordinate environmental sensing with transcriptional control. Future efforts aimed at visualizing CbrXA oligomeric states, reconstituting functional CbrA in defined membrane environments, and probing conformational and ligand binding dependent coupling between CbrA domains will be essential to fully elucidate how transport, signaling, and metabolism are integrated within this unusually sophisticated regulatory protein.

## Materials and Methods

### Protein expression and purification

The overlapped genes encoding *cbrX* and *cbrA* from *P. putida* KT2440 were codon optimized for *E. coli* expression and cloned into a pET28A vector with a C-terminal 8xHIS tag by TWIST Bioscience (San Francisco, CA). The individual *cbrX* and *cbrA-8xHIS* genes were PCR amplified from this vector individually and assembled into a pETDUET-1 vector under the control of individual T7 promoters using NEB HiFi DNA Assembly according to manufacturer’s protocols. The truncated CbrXA construct containing only CbrX and the CbrA SLC5-STAC domains was generated using the NEB Q5 Mutagenesis kit according to manufacturer protocols. The resulting expression plasmids were sequence verified by whole plasmid sequencing at Plasmidsaurus (San Francisco, CA) before being transformed into chemically competent C41(DE3) *E. coli*. A single colony was grown overnight in 100 mL of Lennox Broth (LB) media with 100 *μ*g/mL carbenicillin at 37 °C with shaking at 250 RPM. The starter culture was used to inoculate 4 L of Terrific Broth (TB) media with 100 *μ*g/mL carbenicillin and several drops of antifoam-204 in 3 L baffled Fernbach flasks. Cultures were grown at 37 °C with shaking at 180 RPM to an OD600 of 0.8 before reducing the temperature to 16 °C and inducing protein expression with 1mM Isopropyl thiogalactoside (IPTG). Protein expression was allowed to proceed overnight before harvesting cultures via centrifugation. Bacterial pellets were flash frozen in liquid nitrogen and stored at −80 °C.

For purification bacterial pellets were thawed and resuspended with a Dounce homogenizer in lysis buffer (50 mM HEPES (pH 8), 300 mM KCl, 5 mM *β*-mercaptoethanol, 10% (w/v) glycerol, 2 mM MgCl_2_) with protease inhibitors (1 *μ*g/mL pepstatin A, 1 *μ*g/mL leupeptin, 1 *μ*g/mL aprotinin, 0.6 mM benzamidine) and ~5 *μ*g of recombinantly produced *Serratia marcescens* extracellular nuclease. The resuspended cell slurry was then lysed via sonication on ice, and the lysate was centrifuged at 4000 xg for 20 minutes to remove large debris and unbroken cells followed by ultracentrifugation at 100,000 xg to isolate the membrane fraction. Membranes were resuspended in a Dounce homogenizer on ice in lysis buffer with protease inhibitors and solubilized by adding LMNG detergent to a final concentration of 1% (w/v) and stirring at 4 °C for 1 hour. Insoluble debris was removed by ultracentrifugation at 100,000 xg for 1 hour before proceeding with two-step affinity and size-exclusion chromatography.

Detergent solubilized CbrXA complexes were purified by applying the solubilized membrane fraction to 1 mL Co^3+^ TALON affinity resin in a gravity flow column format. The resin was subsequently washed in buffer A (25 mM HEPES (pH 8), 150 mM KCl, 5 mM *β*-mercaptoethanol, 10% (w/v) glycerol, 0.005% LMNG, and 20 mM imidazole) to remove contaminants. CbrXA was then eluted in buffer A containing 250 mM imidazole before being concentrated in a 100 kDa MWCO centrifugal concentrator. The concentrated protein was then injected on to a Superdex 200 Increase 10/300GL column in 25 mM HEPES (pH 8), 150 mM KCl, 5 mM *β*-mercaptoethanol, and 0.005% LMNG. Peak fractions were pooled and concentrated to ~6 mg/mL in a 100 kDa MWCO centrifugal concentrator for subsequent cryo-EM experiments. For analysis of co-purified histidine with mass-spectrometry CbrXA SLC5-STAC constructs were purified in the same fashion, except the histidine tag was replaced with a twin-STREP tag, captured on streptactin-XT affinity gel, and eluted with 50 mM biotin. Purity of the final samples was assessed by SDS-PAGE analysis and staining with Aquastain (Bulldog Bio, Portsmouth, NH).

### Cryo-EM Imaging and Data Processing

Purified CbrXA constructs were vitrified by applying 3 *μ*L of purified protein to Quantifoil R1.2/1.3 200 mesh Cu grids that had been glow-discharged in a Pelco EasyGlow for 45 seconds at 15mA. Grids were plunge frozen by blotting for 4 seconds on a Vitrobot Mark IV set to 100% relative humidity (RH), 4 °C, and a blot force of 1. Blotted grids were plunge frozen in liquid ethane cooled by liquid nitrogen. An initial screening dataset was collected on a Talos Arctica 200 kV microscope equipped with a Selectris energy filter and Falcon IVi detector, and the final high-resolution dataset of the truncated CbrXA construct was collected on a Titan Krios G4i equipped with a cold-FEG, fringe-free illumination, and a Selectris-X energy filter with a 10 eV slit width. Movies were collected in .EER format using beam-image shift acquisition targeting ~59 holes per stage move. Movies were collected with a pixel size of 0.731 Å/pixel, a total dose of 45 e^−^/Å^2^, and a defocus range of −0.5 to −1.5*μ*m (**Suppl. Table S1**).

All cryo-EM image processing steps were performed in CryoSPARC(35). Raw .EER movies without upsampling were corrected for beam induced motion and the contrast transfer function (CTF) using patch-motion correction and patch-CTF correction. Blob-based particle picking was performed on denoised micrographs, and 2D classification was used to generate 2D averages for template-based picking. Extracted particles were subjected to multiple rounds of 2D classification followed by *Ab-initio* reconstruction, heterogeneous refinement, and non-uniform refinement in CryoSPARC. The final resolution of cryo-EM maps was determined from Fourier Shell Correlation (FSC) of independently refined half-maps using a Gold-standard cutoff of 0.143(36). Data collection and refinement statistics are reported in **Supplemental Table S1**.

### Atomic model building and refinement

The atomic model for CbrXA SLC5-STAC domains was built by first rigid body docking an AlphaFold(37, 38) model of the CbrXA complex generated with localColabFold(39) into the cryo-EM map using UCSF Chimera(40). The model was manually adjusted in COOT(41) to properly fit the cryo-EM density, and refined in real-space using *phenix.real_space_refine* in the PHENIX software suite(42). Iterative rounds of real-space refinement in PHENIX and manual model adjustment in COOT were performed to optimize model geometry and fit to the experimental cryo-EM map. All models were assessed for appropriate stereochemical properties and fit to the experimental cryo-EM map using MolProbity(43) as implemented in PHENIX (**Suppl. Table S1**). Figures of cryo-EM maps and atomic models were created using UCSF Chimera or UCSF ChimeraX(44).

### P. putida strain construction and bacterial growth assays

Knockout strains (Δ*cbrXAB*) of *P. putida* KT2440 were generated by allelic exchange(27). DNA fragments encoding ~750 base pairs upstream and downstream of *cbrXAB* were PCR amplified from *P. putida* KT2440 genomic DNA and assembled into a pEXG2 vector(45). Deletion constructs were designed to maintain the start codon of *cbrX* and ~42 nucleotides at the 3’ end of *cbrB*. This cloning strategy removes the bulk of *cbrXAB* and instead produces a short (~14 amino acid) peptide to avoid potential ectopic effects to neighboring open reading frames. The resulting plasmids were transformed into *P. putida* KT2440 using electroporation, and merodiploids were selected by plating on LB-agar plates containing 30 *μ*g/mL gentamycin and confirmed by colony PCR with primers upstream and downstream of the *cbrXAB* operon. Confirmed merodiploids were then counter-selected by plating on LB-agar plates with 15% (wt/vol) sucrose. Sucrose insensitive colonies were screened for deletion of *cbrXAB* by colony PCR and further confirmed by nanopore sequencing at Plasmidsaurus.

To complement knockout strains of *P. putida* the wild-type *cbrXAB* sequence including the upstream P_cbrA_ promoter was cloned into a pUC18T-mini-Tn*7*T-Gm vector(28) using HiFi DNA Assembly. The resulting plasmid was then used as a template to make domain deletion constructs using the NEB Q5 Mutagenesis Kit according to manufacturer protocols. The resulting plasmids along with a pTNS2 helper plasmid were transformed into the Δ*cbrXAB* strains by electroporation and plated on LB-agar containing 30 *μ*g/mL gentamycin(28). Transformants that had integrated *cbrXAB* at the mini-TN7 site downstream of *glmS* were screened by colony PCR followed by nanopore sequencing at Plasmidsaurus.

Bacterial growth assays were performed by growing a single colony of each *P. putida* strain in 3 mL LB medium overnight at 30 °C with shaking at 250 RPM. The following morning 1 mL of culture was pelleted by centrifugation and resuspended in 7 mL PBS before being diluted to a final OD600 of 0.05 in M9 minimal medium (42.2 mM Na_2_HPO_4_, 22 mM KH_2_PO_4_, 8.6 mM NaCl, 2 mM MgSO_4_, trace elements) containing L-histidine (20 mM) as a sole carbon and nitrogen source. Cells were plated in clear 96-well plates (200 *μ*L/well) and growth at 30 °C was monitored at OD600 in a Molecular Devices iD5 plate reader with read intervals of 10 minutes with continuous medium intensity double orbital shaking. Growth assays were performed in biological triplicate.

### Molecular dynamics simulations and analysis

The structure of CbrXA built from cryo-EM data was embedded in a POPC bilayer using CHARMM-GUI membrane builder(46). The obtained membrane-protein systems were neutralized, and NaCl was added to achieve a salt concentration of 150 mM and solvated with TIP3P water molecules extending at least 20Å on each side. The final systems containing ~205,000 atoms with the box size (132 Å × 132 Å × 126 Å), were made periodic on all three directions and periodic boundary conditions were applied. Another system with deprotonated K196 was also prepared identically to investigate the impact of protonation state on conformational space of CbrXA and corresponding histidine motion. All simulations were performed using OpenMM 8(47) with the CHARMM36m forcefield(48) using the Langevin integrator(49) with 2 fs timestep. The systems were first energy minimized to relax the protein and lipids and remove steric clashes. The minimized systems were subjected to NVT equilibrations, followed by NPT equilibrations and MD production in NPT ensemble at 310 K and 1 atmospheric pressure using Monte Carlo barostat(50). The force-based switching method was used for smoothing vdW interactions over the distance range 10-12 Å, and the particle-mesh Ewald (PME) method(51) was used for long-range electrostatic interactions. Five independent replicates were simulated for each protonation state ranging between 900-1100 ns with different initial velocities to ensure reproducibility and adequate sampling of the conformational dynamics. The simulation trajectories were visualized using VMD(52) and molecular graphics in the figures were rendered in both VMD and UCSF Chimera X(44). Water occupancy maps were generated by combining trajectories from all simulations and analyzing them with the VolMap plugin in VMD. Trajectory frames after 400 ns were included, since we notice a steady state in water permeation i.e., a plateau in water count around the substrate and the break region, around that time-point in our simulation. Water occupancy within 10 Å of the substrate histidine or CbrA residue K196 was quantified by computing an average occupancy grid at an isosurface threshold of 0.1, corresponding to water being present in a voxel in 10% of snapshots. The resulting occupancy maps were overlaid onto the cryo-EM–derived atomic coordinates to compare experimentally observed water positions with those identified in the MD simulations.

Subsequent analyses such as (i) computing root mean square deviations (RMSD) of substrate histidine and break region residues, (ii) building contact maps of histidine with CbrA along the trajectories and (iii) carrying out time-lagged independent component analysis (tICA) to extract slow motions from the MD ensembles were performed using scripts from the MD_Interpret, MDtraj libraries(53) and Deeptime packages(54). To remove global translational and rotational motion, the deprotonated and protonated MD trajectories were mean-centered and aligned to their respective post-equilibration reference frames. The backbone atoms were used as the alignment indices for all trajectory superpositions. It should be noted that there is negligible difference between the post-equilibration structures of deprotonated and protonated system (backbone RMSD < 0.2 nm). The feature sets to train tICA models were built using either the aligned histidine substrate coordinates or the aligned TM2 break region heavy atoms. The tICA method was used to identify the slow collective motions underlying the MD trajectories and to construct a reduced representation of the conformational landscape (55). In this framework, structural descriptors of the system (the aligned coordinates of histidine or break region residues) are linearly transformed into a set of orthogonal coordinates that maximize time-lagged autocorrelation at a chosen lag time. This procedure provides an approximation to the slow eigenmodes of the underlying dynamical propagator that governs the time evolution of the molecular system. By emphasizing motions that relax slowly relative to the lag time, tICA filters out fast fluctuations and isolates collective coordinates associated with transitions between metastable conformational states. The leading time- lagged component(s) therefore provide a kinetically meaningful low-dimensional coordinate system for analyzing conformational transitions in the simulation trajectories (34). In this work, we examined a range of model parameters, including lag times from 10 ns to 200 ns and tICA dimensionalities between 2 and 5. Across this range, the qualitative structure of the free-energy landscape projected onto the leading time-lagged independent components (tIC1 and tIC2) remained largely unchanged, indicating that the dominant slow dynamical modes captured by the tICA models are robust to the choice of these parameters. We have used a sufficiently large lag time of 100 ns and two component tICA model to present our free energy analysis in the Results section, with similar free energy distributions using different model parameters are provided in the SI.

### Statistical Analysis

*P. putida* growth assays were performed in biological triplicate (N=3) by starting growth protocols from independently picked individual bacterial colonies. Biological replicates were also measured in technical triplicate, and the reported values represent the average of values obtained across biological replicates (measured in technical triplicate). Where appropriate, error bars represent the standard error of the mean (SEM) across biological replicates.

## Acknowledgments

Electron microscopy data was collected at the RTSF Cryo-Electron Microscopy Core at Michigan State University. We are grateful to Dr. Sundharraman Subramanian for assistance with cryo-EM data collection. The pEXG2 vector was a generous gift from Dr. Albert Siryaporn from the University of California Irvine. We are also grateful to Dr. Alex Dickson, Bradon Krah, Peixuan Yu, and Dr. Sundharraman Subramanian for review of the manuscript and thoughtful discussions.

## Author Contributions

MAO and MMF performed molecular cloning and protein purification. MAO constructed bacterial strains and performed growth assays and analysis. MMF and BJO collected and processed cryo-EM data. BJO built and refined the atomic model. TS and SB performed molecular simulations and data analysis. SB oversaw molecular simulations. BJO conceived the project, and wrote the manuscript with input and refinement from all authors.

## Data and materials availability

Atomic coordinates and associated electron microscopy maps for the structure reported in this publication have been deposited in the Protein Data Bank (PDB) and Electron Microscopy Data Bank (EMDB) under the following accession number: CbrXA SLC5-STAC (PDB 10ED, EMDB 75104). All files required to visualize and reproduce molecular dynamics simulations are freely available on Zenodo under the following DOI (10.5281/zenodo.18867874). The analysis codes for RMSD, tICA, and water permeation are available on Github (https://github.com/SamikBose/MD_Interpret). All requests for plasmids, bacterial strains, or other materials will be honored upon request to BJO with an appropriate Materials Transfer Agreement issued by Michigan State University.

## Supplementary Materials

Supplementary materials including Supplemental Figures S1-S9 and Supplemental Table S1 are provided with this manuscript.

