## Supplemental Data for "Structure and conformational dynamics of the *Pseudomonas* CbrA transceptor"

**This PDF file includes:**

Figures S1 to S9
Table S1

### Figures

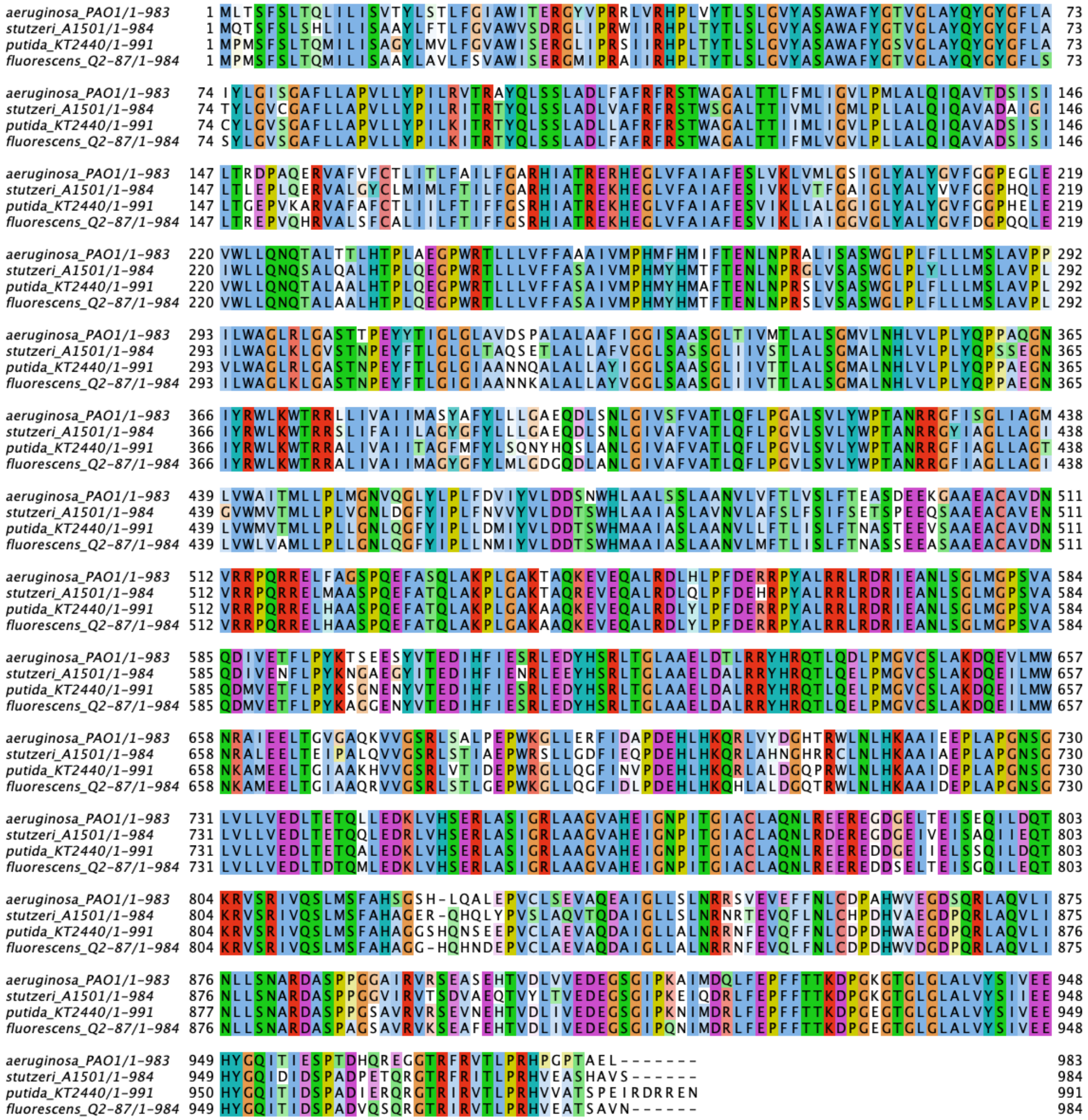

**Fig. S1. Sequence alignment of CbrA proteins**

Shown is a sequence alignment of CbrA protein from four different strains of *Pseudomonas*. Strain names with protein length are indicated to the left. Sequence alignment was performed in Clustal Omega with visualization using Clustal coloring as implemented in JalView. *P. putida* CbrA is 82% identical to the protein from *P. aeruginosa.*

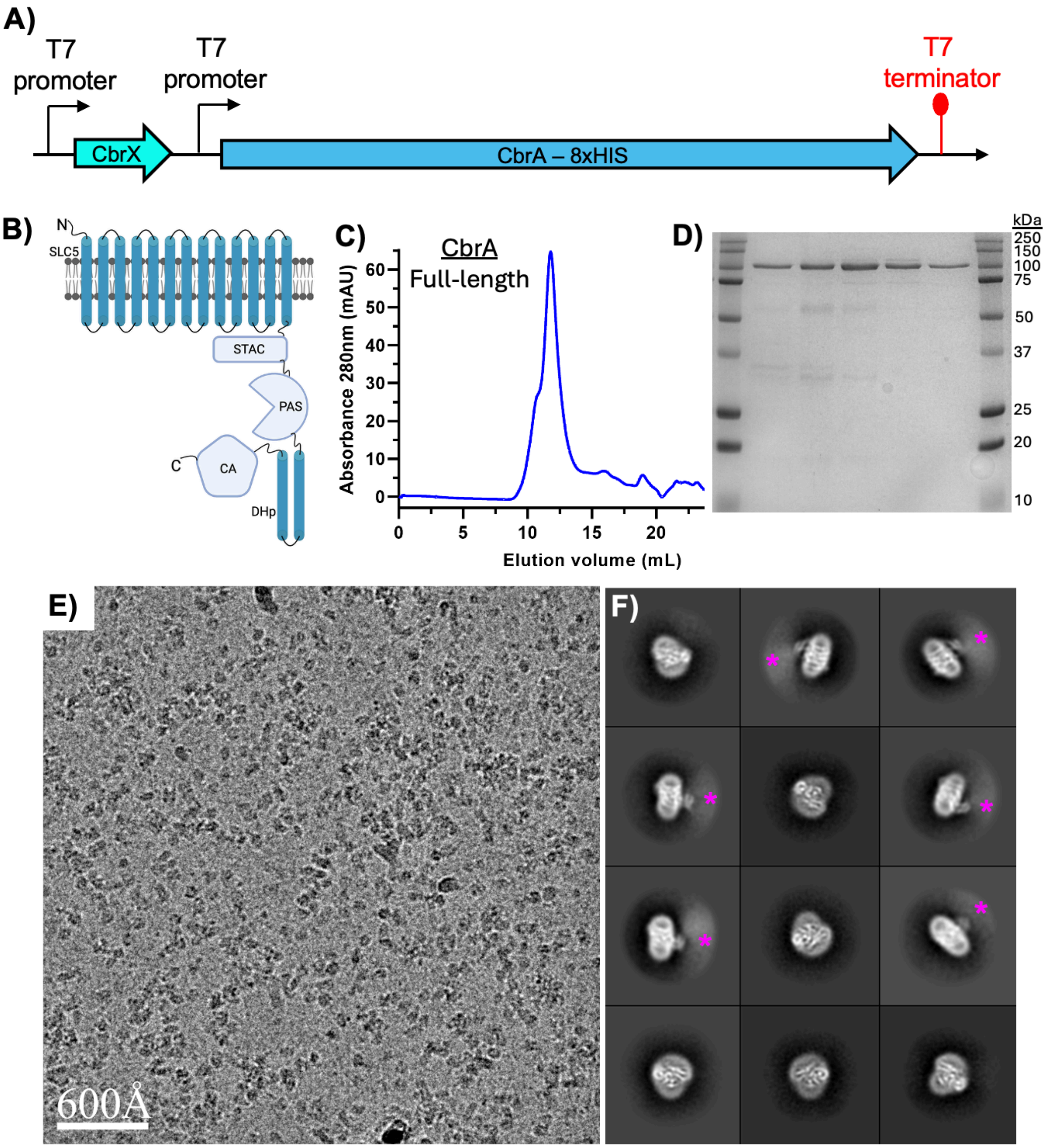


**Fig. S2. Purification and cryo-EM analysis of full-length CbrXA**
**A**) Schematic showing the petDUET expression construct for CbrXA. CbrX and CbrA were separated with expression driven from individual T7 promoters. **B**) Schematic of a full-length CbrA monomer with individual domains labeled. Full-length CbrA has a molecular weight of ~109 kDa. **C**) Gel-filtration chromatogram of full-length CbrXA purified in detergent. **D**) SDS-PAGE analysis of peak fractions from the Gel-filtration chromatogram shown in B. **E**) Electron micrograph of full-length CbrXA particles acquired on a 200 KeV Talos Arctica. Micrograph is low pass filtered to 5Å to facilitate particle visualization. **F**) 2D class averages of full-length CbrXA particles. The averages demonstrate that the soluble domains of CbrA (PAS, DHp, CA) are disordered and not visible (pink asterisks).

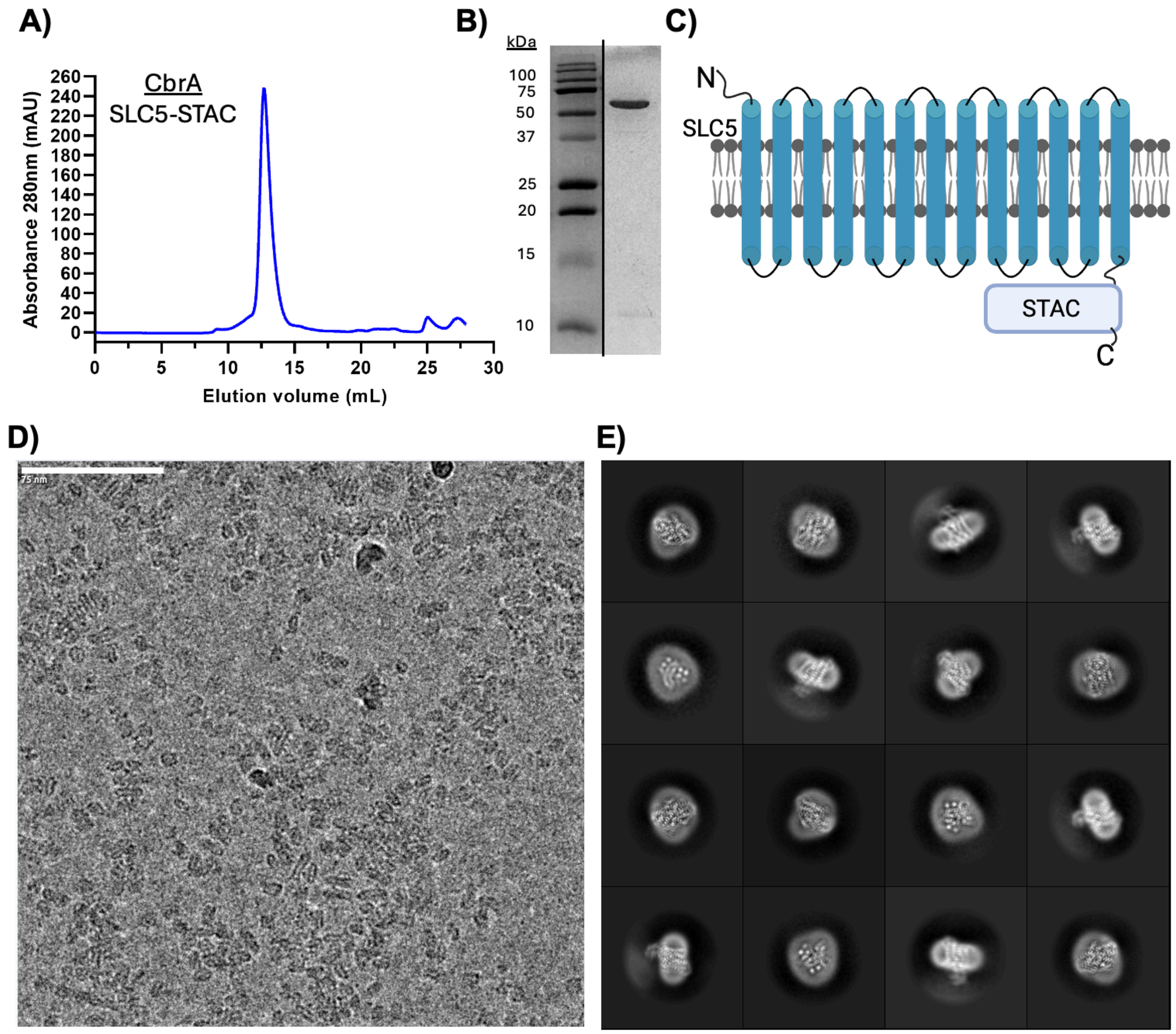


**Figure S3. Structural analysis of truncated CbrXA**
**A)** Gel-filtration chromatogram of truncated CbrXA purified in detergent. **B)** SDS-PAGE analysis of peak fraction from the chromatogram shown in **A**. **C)** **S**chematic of the truncated construct with PAS, DHp, and CA domains removed. Truncated CbrXA has a molecular weight of ~66 kDa. **D)** Electron micrograph of CbrXA SLC5-STAC particles acquired on a 300 KeV Titan Krios. Micrograph is low pass filtered to 5Å to facilitate particle visualization. **E)** 2D class averages of CbrXA SLC5-STAC particles

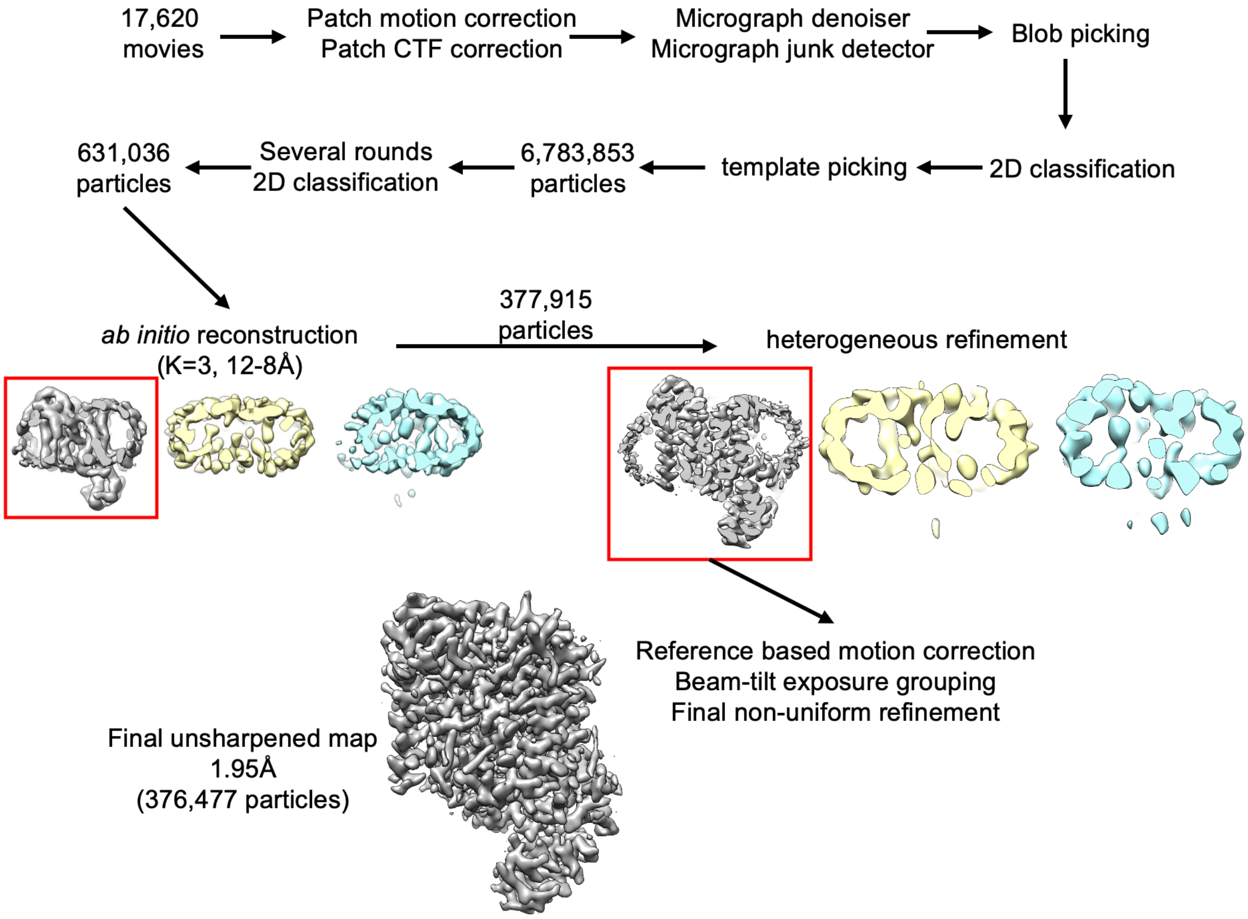


**Figure S4. Cryo-EM data processing for truncated CbrXA**
**A)** Data processing workflow for truncated CbrXA SLC-STAC reconstruction. All steps were performed in CryoSPARC. Red boxes indicate classes that were selected for processing in subsequent steps.

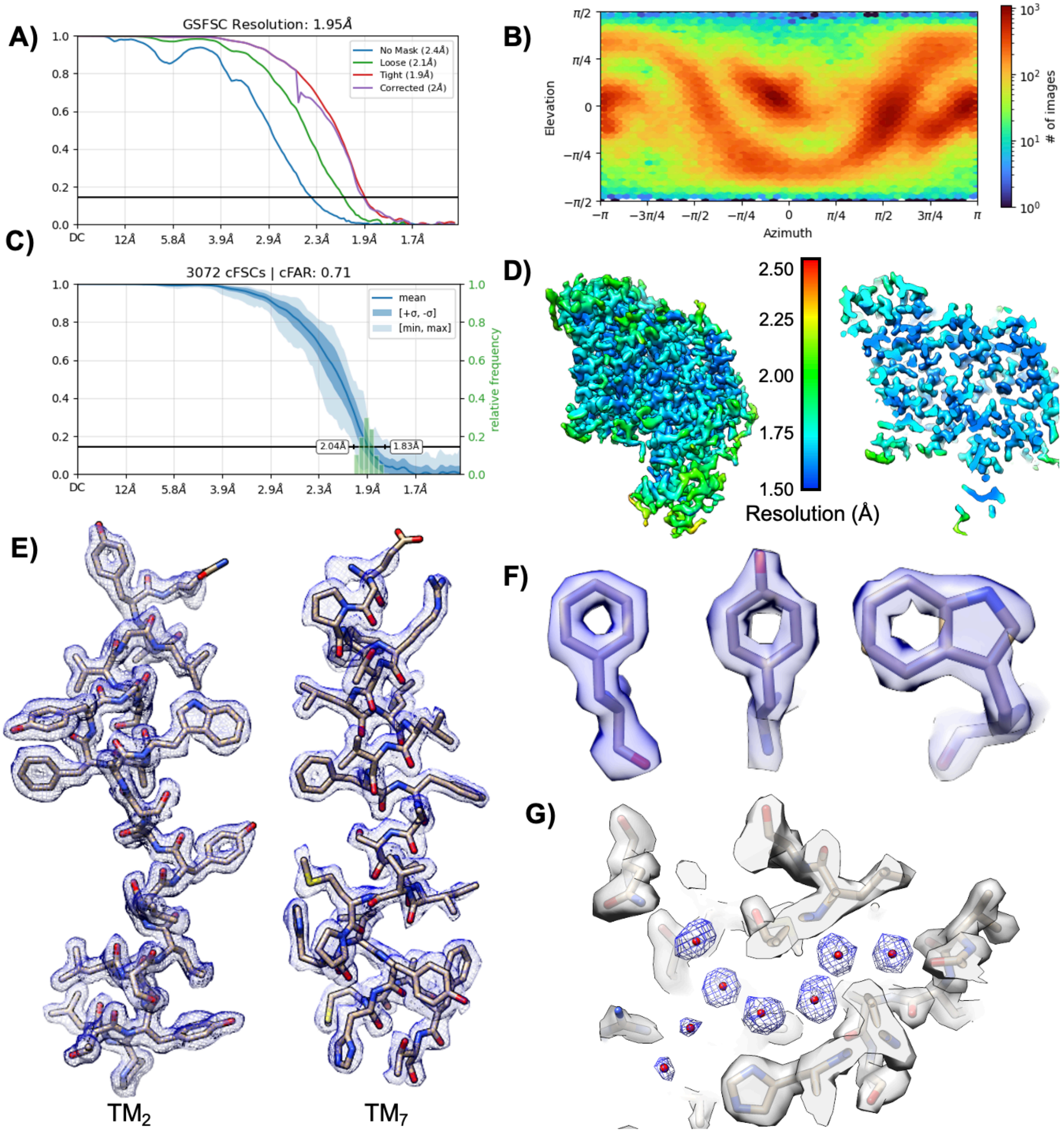


**Figure S5. Cryo-EM data analysis for CbrXA SLC5-STAC construct**

**A)** Gold-standard Fourier shell correlation (FSC) curve of final 3D reconstruction. **B)** Angle distribution of particles in the final 3D reconstruction. **C)** Directional FSC curve of the final reconstruction. **D)** Local resolution map of final 3D reconstruction. An overall view (left) and sliced view (right) are provided. Most regions of the TM helices are resolved to better than 1.8Å. **E)** Representative coulomb potential map for two transmembrane helices demonstrating clear resolution of sidechain rotamers and backbone carbonyls. **F)** Coulomb potential map for selected sidechains demonstrating sufficient resolution to resolve clear holes for aromatic residues. **G)** Densities for waters (blue mesh) are visible throughout the reconstructed map.

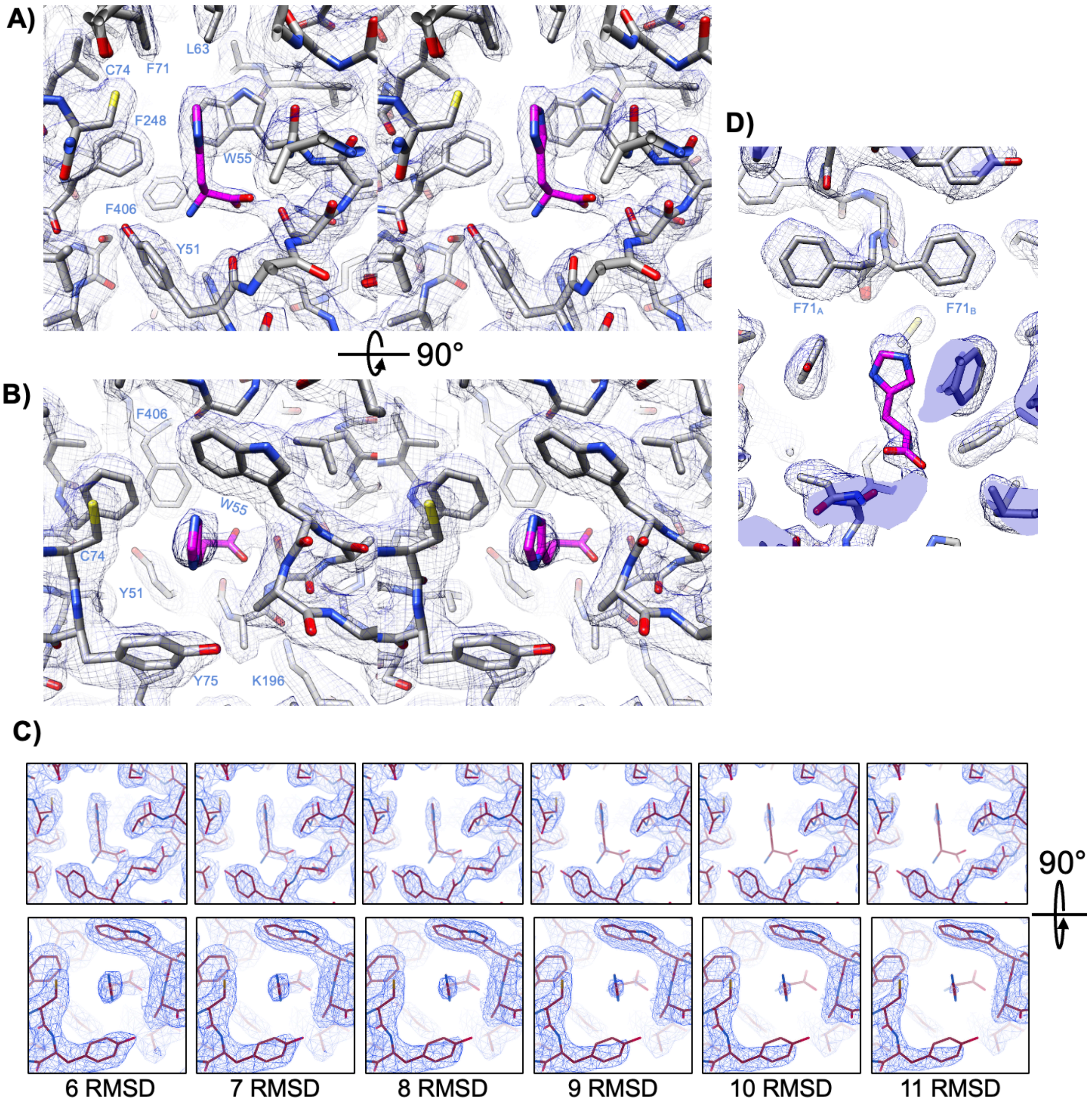


**Figure S6. CbrA histidine binding pocket**

**A&B)** Stereo diagram of the histidine binding pocket in the CbrA SLC5 domain. Histidine is colored magenta, and CbrA residues are colored grey. The unsharpened coulomb potential map is shown in blue mesh. Panel **B** is rotated 90° towards the viewer compared to panel **A**, and provides a view from the periplasmic space. **C)** View of the coulomb potential map in the histidine binding pocket at different map contour levels. The histidine is centered in each image. Density for the co-purified histidine is weaker than surrounding residues of CbrA, suggesting possible low occupancy of histidine in the pocket. **D)** View of alternate rotamer state modeling of residue F71. Alternate rotamers are labeled $F71_A$ and $F71_B$.

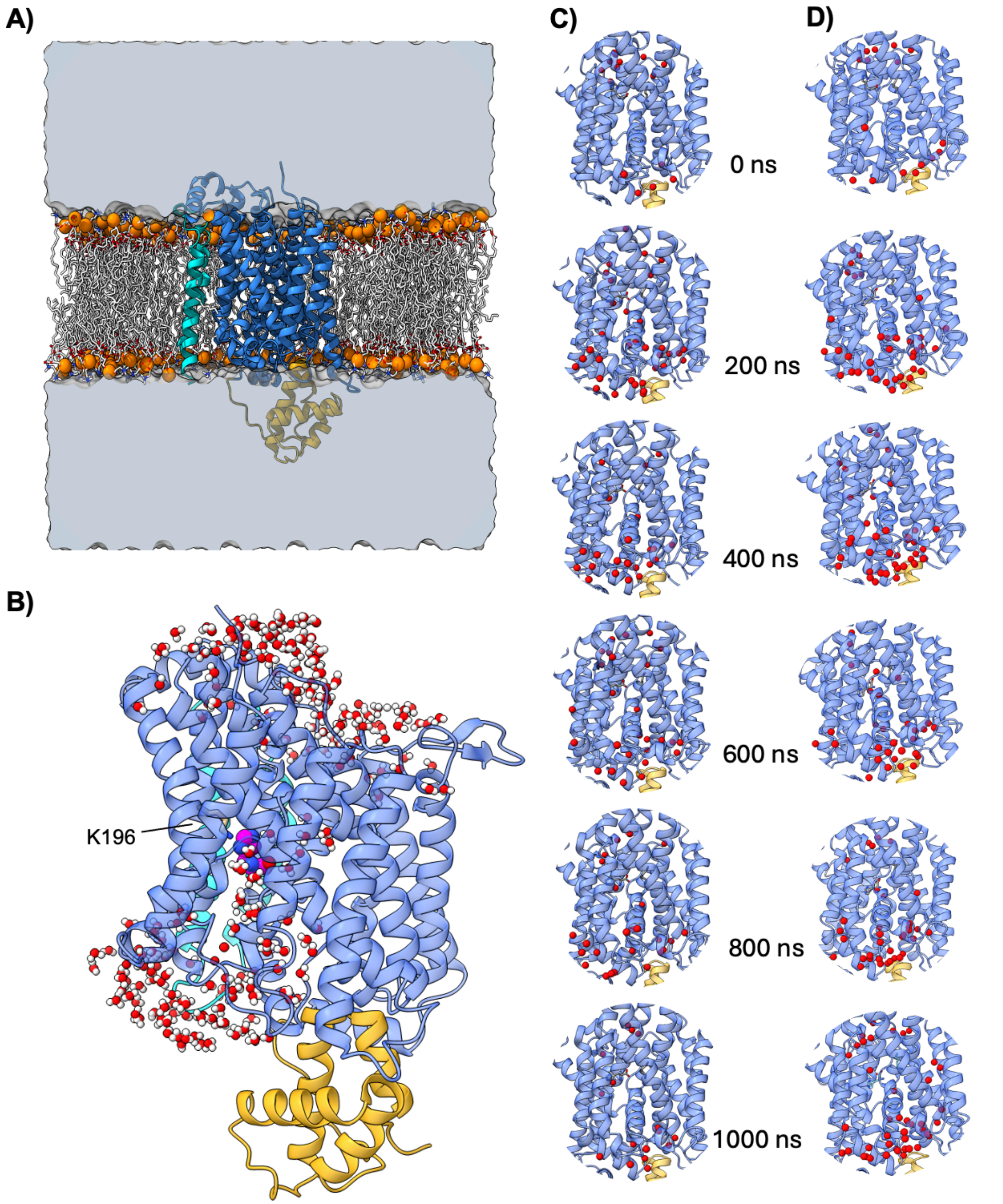


**S7. Molecular dynamics simulations**

**A)** View of the MD simulation box prior to equilibration showing CbrXA embedded in a lipid bilayer with surrounding solvent. CbrXA is shown as a cartoon and colored the same as in main figures. Lipid phosphate groups are shown as orange spheres. Bulk solvent on either side of the membrane is shown as a grey surface. **B)** View showing hydrogen-bonded water network extending from cytosolic region to the vicinity of K196. **C&D)** Water permeation through the SLC-STAC domain in protonated (**C**) and deprotonated (**D**) simulations. SLC5 domain is shown in blue, STAC domain in orange, and waters are shown in red spheres.

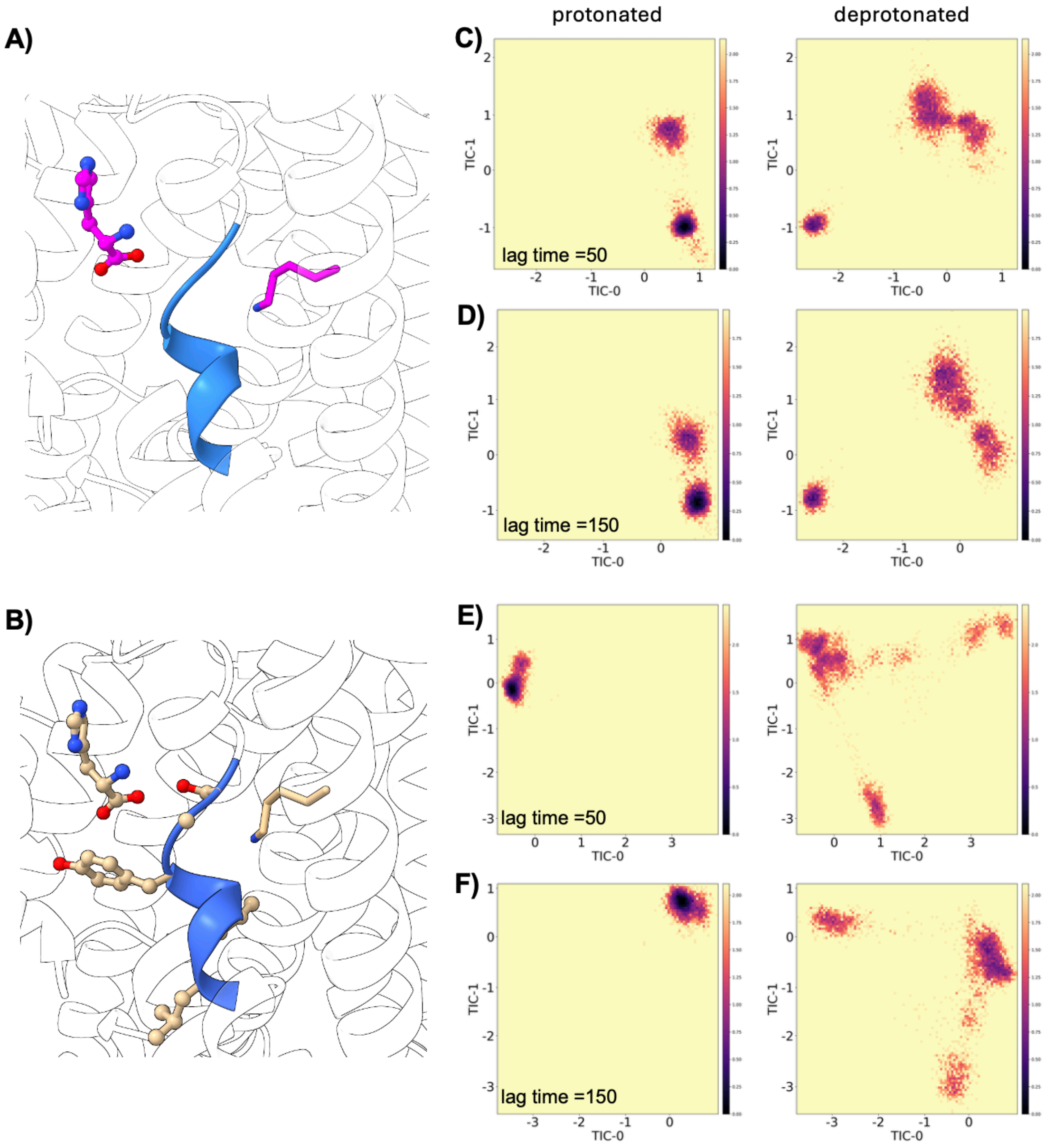


**Figure S8. tICA features and analysis**
Features used for tICA training **A)** histidine heavy atoms, shown as ball-and-stick models in magenta, and **B)** heavy atoms of the break region residues, shown in beige. Histidine and K196 are also displayed to show the relative positions of these residues. tICA plots using histidine backbone features at lag times of **C)** 50 and **D)** 150 are shown for the protonated (left) and deprotonated (right) systems. Similarly, plots **E)** and **F)** show the tICA projections using CbrA backbone residues as features at lag times of 50 and 150, respectively.

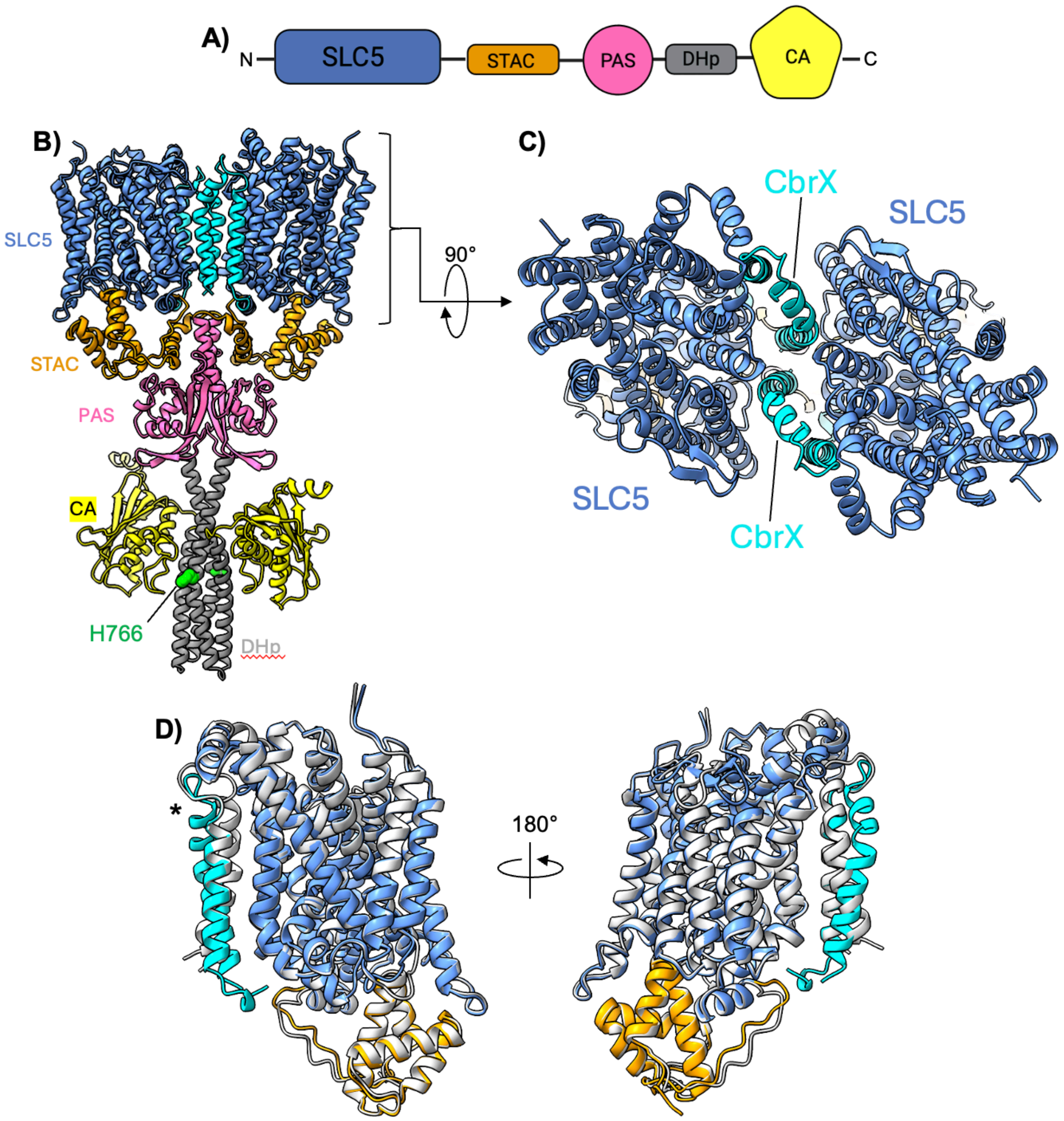


**Figure S9. Alphafold models of CbrXA**

**A)** Domain organization of CbrA. Amino and carboxy termini are labeled. **B)** Alphafold-3 model of CbrXA dimer. CbrA domains are colored as shown in **A**, and CbrX is colored in cyan. Histidine 766 that is phosphorylated by the CA domain is colored green. **C)** Rotated view of the membrane region shown in **B**. Two CbrX peptides form the dimer interface between CbrA SLC5 domains. **D)** Comparison of Alphafold predicted CbrXA structure and that determined by cryo-EM. The Alphafold model is shown with coloring consistent with panel **A**, the cryo-EM derived model is shown in grey. Overall RMSD between $C_{\alpha}$ carbons is 1.4Å. The largest deviation is in the extracellular tip of CbrX (marked with an asterisk).

**Tables**

| | CbrXA SLC5-STAC (PDB: 10ED ) (EMDB: 75104) |
|---|---|
| **Data collection and processing** | |
| Magnification | 165,000 |
| Voltage (kV) | 300 |
| Electron exposure (e-/Å) | 45 |
| Defocus range ($\mu$m) | -0.5 to -1.5 |
| Pixel size (Å) | 0.731 |
| Symmetry imposed | C1 |
| Number of micrographs (#) | 17,629 |
| Map resolution (Å) | 1.95 |
| FSC threshold | 0.143 |
| **Refinement** | |
| Model resolution | 1.99 |
| FSC threshold | 0.5 |
| Map sharpening *B* factor (Å$^2$) | -48.0 |
| Model composition | |
| Non-hydrogen atoms | 5448 |
| Protein residues | 647 |
| Ligands | 18 |
| Mean *B* factors (Å$^2$) | |
| Protein | 27.45 |
| Ligand | 46.28 |
| R.m.s.deviations | |
| Bond lengths (Å) | 0.003 |
| Bond angles (°) | 0.564 |
| Validation | |
| MolProbity score | 1.25 |
| Clashscore | 4.77 |
| Poor rotamers (%) | 0.76 |
| Ramachandran plot | |
| Favored (%) | 99.07 |
| Allowed (%) | 0.93 |
| Disallowed (%) | 0.00 |

**Table S1. Cryo-EM Data Collection and Refinement Statistics**
